# Behavioral expression of decision confidence engages disentangled confidence coding in the orbitofrontal cortex

**DOI:** 10.1101/2025.08.29.673022

**Authors:** Tomoya Ohnuki, Yuma Osako, Kazuki Shiotani, Yuta Tanisumi, Shogo Takamiya, Hiroyuki Manabe, Yoshio Sakurai, Junya Hirokawa

## Abstract

Task engagement is thought to recruit subsets of neurons encoding task-relevant variables, but it remains unclear whether it also reorganizes the geometry of population codes and how such reorganization supports behavior. Here we show that, in the orbitofrontal cortex (OFC), task engagement aligns population activity along a confidence axis, thereby enabling the behavioral use of decision confidence. We recorded OFC activity while rats performed two variants of the same perceptual decision-making task that differed only in reward-timing uncertainty, which altered the utility of post-decision confidence for guiding reward waiting. OFC neurons exhibited more linear responses to confidence when confidence was more strongly expressed in behavior. Although the proportion of OFC neurons encoding confidence was similar across strategies, strategy-dependent alignment of population activity to confidence emerged under variable long delays, and this alignment predicted confidence-based waiting behavior. These findings suggest that OFC population codes flexibly adapt to the behavioral relevance of task variables, linking cognitive strategy to the geometry of neural representation.

## Introduction

The engagement of cognitive functions is thought to recruit neurons that represent information relevant to the task at hand. A long-standing benchmark for understanding how brain regions support cognition has been the identification of neurons whose activity reflects such task-relevant information. Traditionally, the number of such neurons has been considered evidence for the involvement and functional significance of a given brain region in supporting cognitive abilities expressed through behavior. For example, numerous studies have reported the emergence and strengthening of neuronal representations—such as firing selectivity to sensory stimuli or motor responses associated with rewarding and aversive outcomes—across various brain regions as animals learn cognitive tasks^1–8^. Moreover, numerous studies have demonstrated that the level of task engagement— whether animals are actively or passively involved—can modulate the recruitment of neurons encoding task-relevant variables, even after learning has been established^9–14^. Although these findings suggest a close relationship between neuronal recruitment and the behavioral expression of cognitive functions, most of them do not clearly dissociate neural activity reflecting cognitive processes from activity reflecting concurrent behavioral changes.

One potential advantage of recruiting large numbers of neurons is the capacity for population coding, in which information is distributed across neurons^15–17^. Population coding has been shown to confer several computational and representational benefits, including enhanced robustness to noise^18,19^, the ability to maintain information over time^20–22^, and the flexibility to support linearly decodable and disentangled representations of abstract variables^23–28^. These features align with key hallmarks of higher-order cognition, many aspects of which are associated with the frontal cortex. From this perspective, the extent and structure of neuronal recruitment may reflect the degree to which specific cognitive functions—particularly higher-order ones—are engaged. However, no studies have directly tested whether engaging a higher-order cognitive strategy improves performance by simply recruiting more neurons, or by restructuring the population code of those neurons.

To address this gap, we designed an experiment that isolates cognitive strategy from motor output and reward contingencies by holding the latter constant. Specifically, we examined how neurons in the orbitofrontal cortex (OFC)—a region previously shown to encode post-decision confidence and to play a causal role in its behavioral expression—represent task-relevant information during a deliberative decision-making task.^29–33^. By manipulating the temporal uncertainty of reward delivery, we induced the same animals to flexibly engage or disengage confidence-based strategies depending on task demands. Interestingly, the proportion of OFC neurons encoding post-decision confidence was comparable across task conditions. However, when confidence was overtly expressed in behavior, its neural representation became more temporally stable and more linearly decodable across a continuum of confidence levels. These findings suggest that the linear accessibility of information guiding behavior—rather than the sheer number of neurons representing it— more accurately reflects the expression of higher-order cognitive functions.

## Results

### Rats adopted a confidence-dependent strategy based on the timing uncertainty of potential rewards

We trained rats in a decision-making task, which entailed determining the duration they would wait for potential rewards after making perceptual decisions (Fig. 1A)^30,32^. Each trial began with the rats poking their snouts into the central port to receive olfactory cues. Based on this cue, the rat made a perceptual decision by choosing either the left or right port. Task difficulty was systematically manipulated by presenting binary mixtures of two odorants— each associated with a different choice side—at varying concentration ratios^34^. Only correct choices, defined as those consistent with the dominant odor in the mixture, were rewarded following a delay period. Importantly, no explicit feedback was provided for incorrect choices. Thus, the rats had to decide how long to wait for a potential reward on each trial, with the waiting duration serving as a measure of their time investment in the perceptual decision. To avoid confounding the time spent consuming rewards with reward waiting time in correct trials, we omitted rewards in 20% of trials (probe trials). Rats almost perfectly discriminated the easiest cues, and choice accuracy was strongly correlated with odor difficulty (Fig. 1B), confirming that the odor mixtures effectively modulated decision difficulty (7 rats, 134 sessions, 117,871 trials).

**Fig. 1:**
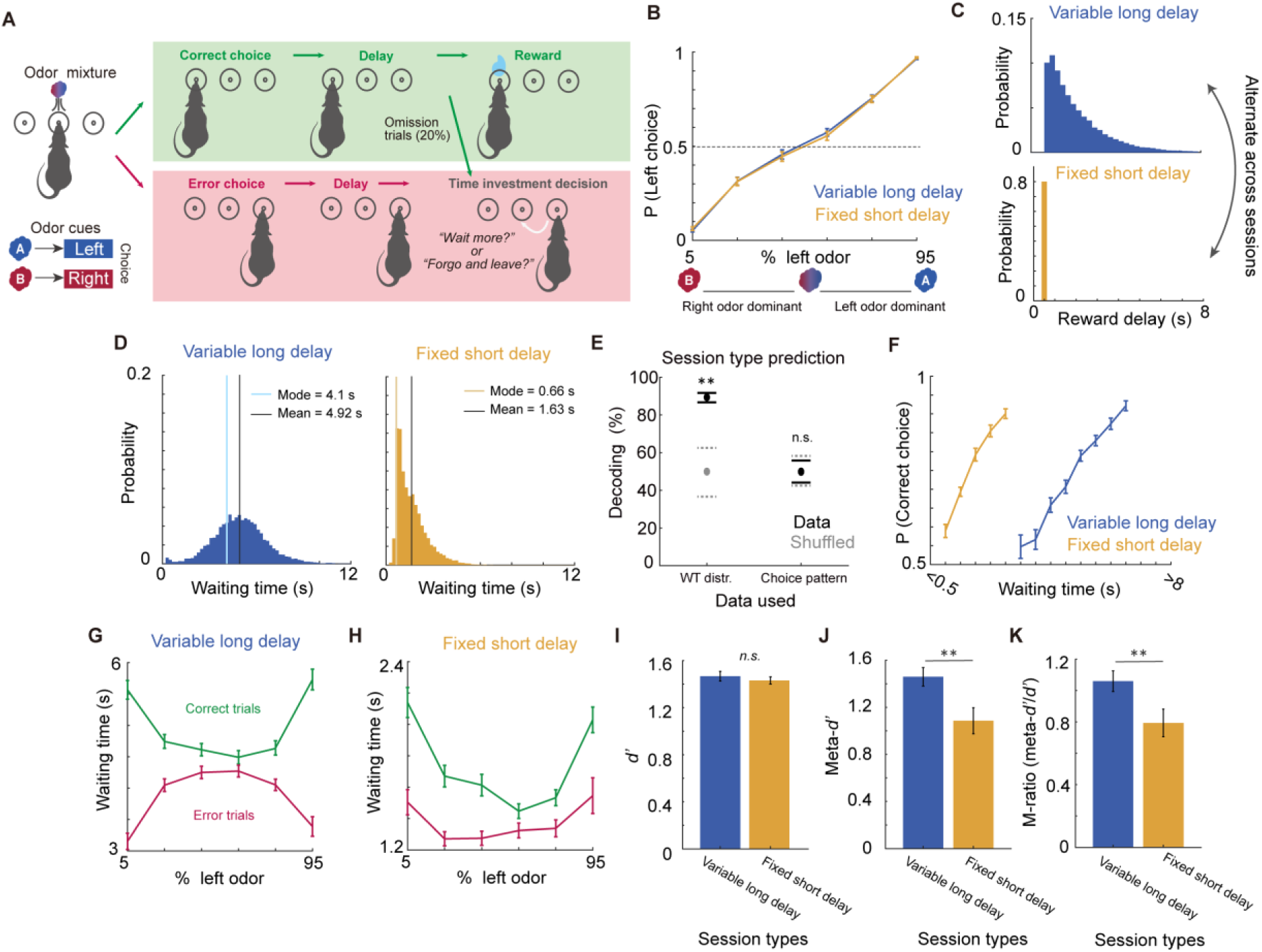
Altered behavioral expression of confidence driven by reward timing uncertainty. (A) Schematic timeline of the behavioral task. (B) Odor discrimination performance across sessions (*n* = 67 sessions per session type from 7 rats). Lines indicate session types (blue: variable long delay; orange: fixed short delay). (C) Example distributions of reward delay used in the variable long delay (top) and fixed short delay (bottom) sessions. (D) Distributions of reward waiting times pooled across all sessions for the variable long delay (left) and fixed short delay (right) conditions. Colored lines indicate the mode (light blue or orange) and the mean (black) of the reward waiting time. (E) Decoding accuracy of session type based on behavioral features. Black circles and lines indicate the mean and 95% range (2.5th to 97.5th percentile) for the real data, compared against models (gray) trained with label-shuffled data. Asterisks indicate statistical significance (bootstrap *n* = 100; estimated *p* < 0.05; see Methods). (F) Correlation between reward waiting time and correct choice rate in each session type. (G) Reward waiting time in the long delay sessions across combinations of sensory evidence and choice outcome. These combinations correspond to different average confidence levels, as predicted by signal detection theory. (H) Same as G, but for the short delay sessions. (I) Average *d’* across different session types (blue: variable long delay; orange: fixed short delay). (J) Same as I, but for meta-*d’*. (K) Same as I and J, but for M-ratio. (B–K) Lines show the mean values, and error bars show the standard error of the mean (SEM). (I-K) Asterisks indicate statistical significance (t-test, *p* < 0.05); *n.s.* indicates non-significance.

The rats alternately performed the task with two types of reward delay distributions across days: variable long or fixed short, differing in the delay length and uncertainty of reward timing (67 sessions each; 54,708 and 63,163 trials, respectively). In a variable long delay session, potential reward delays were drawn from an exponential distribution (0.8-8 s), maintaining a relatively constant reward expectation across delays (top in Fig. 1C)^30,32,35–37^. Conversely, in a fixed short delay session, potential rewards were consistently delivered 0.5 s after the choices, allowing the rats to anticipate the timing of the reward (bottom in Fig. 1C). Notably, the difference in reward timing uncertainty did not impact the perceptual decision strategy, as indicated by comparable choice accuracy, which varied with odor difficulties similarly between the two session types (Fig. 1B). However, the distribution of reward waiting time following decisions showed a prominent shift. While waiting times in the variable long delay session symmetrically varied around the mean, those in the short delay session exhibited a right-skewed distribution with a clear peak around the reward timing of 0.5 s (Fig. 1D). Changes in the waiting time distribution—specifically in skewness and kurtosis, which are independent of the mean—were sufficient to predict session types (Fig. 1E), whereas choice patterns across odor mixtures (Fig. 1B) were not.

The above result suggested that the reward waiting time, time investment in perceptual decision known reflecting post-decision confidence^30,32^, was specifically modulated by the reward timing uncertainty. Notably, in the fixed short delay sessions, the reward waiting time was closely aligned with the expected reward timing (Fig. 1D, right). To investigate the underlying behavioral strategies, we analyzed how reward waiting time correlated with decision confidence, which we operationalized as statistical decision confidence: the probability of a correct choice given the subjective evidence and the choice, *P(choice = correct | subjective evidence, choice)*^29,32,38^. We examined whether the observed behavior exhibited features of the statistically defined confidence, as described in previous studies^32,39^. First, if waiting time reflects confidence, it should correlate with choice accuracy. Consistent with this expectation, we found that choice accuracy increased with waiting time in both session types (Fig. 1F), indicating that rats estimated their own accuracy to guide time investment. Notably, the contrast in accuracy between the lowest and highest waiting-time ranges was greater in the long-delay sessions than in the short-delay sessions, suggesting a more explicit expression of confidence. Next, we asked whether waiting time varied systematically with the interaction between sensory evidence and choice outcome. According to signal detection theory and previous studies^32,39^, confidence is generally expected to increase with sensory evidence strength for correct choices but decrease for incorrect choices. In the long delay sessions, reward waiting time followed this canonical pattern, forming an X-shaped pattern (Fig. 1G), consistent with previous findings. In contrast, this pattern was attenuated in the fixed short delay sessions (Fig. 1H). While waiting times still differed between correct and error trials, error trials exhibited a moderate U-shaped pattern—similar to that of correct trials—suggesting reduced integration of sensory evidence and outcomes, and a diminished use of confidence in guiding time investment behavior under predictable reward timing (Supplementary Fig. 1).

To further support this interpretation, we computed the M-ratio (meta-d’/d’), a widely used measure of the efficiency of metacognitive performance relative to sensory discrimination performance^40,41^. The M-ratio was significantly lower in the short delay sessions than in the long delay sessions (Fig. 1K; t-test, *p* = 0.007), driven specifically by a reduction in meta-*d’* (Fig. 1J; t-test, *p* = 0.004), with no significant difference in *d’* (Fig. 1I; t-test, *p* = 0.316).

Together, these results indicate that rats flexibly adjust their use of post-decision confidence in response to task demands.

### Behavioral impact of confidence is overridden by other task variables when reward timing is highly predictable

Thus far, we have demonstrated that the time investment strategy in perceptual decision-making is modulated by uncertainty of reward timing, as evidenced by multiple behavioral measures. To further investigate how rats adjusted their strategy, we constructed a Generalized Linear Model (GLM) to predict trial-by-trial reward waiting times based on task-related variables. These included decision confidence (modeled as the interaction between sensory evidence and choice outcome as described above), odor difficulty (low, medium, or high), chosen direction (left or right), previous trial outcome (rewarded or unrewarded), and satiety (quantified as the cumulative reward amount up to the current trial) (Fig. 2A–B). The relative contribution of each variable was quantified by measuring the decrease in explained variance (ΔR²) when a given parameter was excluded from the full model (Fig. 2C)^42^. Consistent with the above findings, confidence had the greatest impact on model performance, although its contribution was markedly reduced in the fixed short delay sessions (Long: 66.8 ± 3.2%; Short: 45.6 ± 3.5%). This indicates that, with predictable reward timing, the time investment strategy relied less on confidence. Conversely, the contributions of odor difficulty (Long: 4.5 ± 0.8%; Short: 14.5 ± 1.8%), choice direction (Long: 9.2 ± 1.9%; Short: 22.1 ± 3.3%), and reward satiety (Long: 8.2 ± 1.6%; Short: 14.5 ± 2.4%) were higher in the short delay sessions. These shifts underscore a greater impact of sensory and internal state variables in short delay sessions, coupled with a diminished expression of confidence due to the predictability of immediate rewards.

**Fig. 2:**
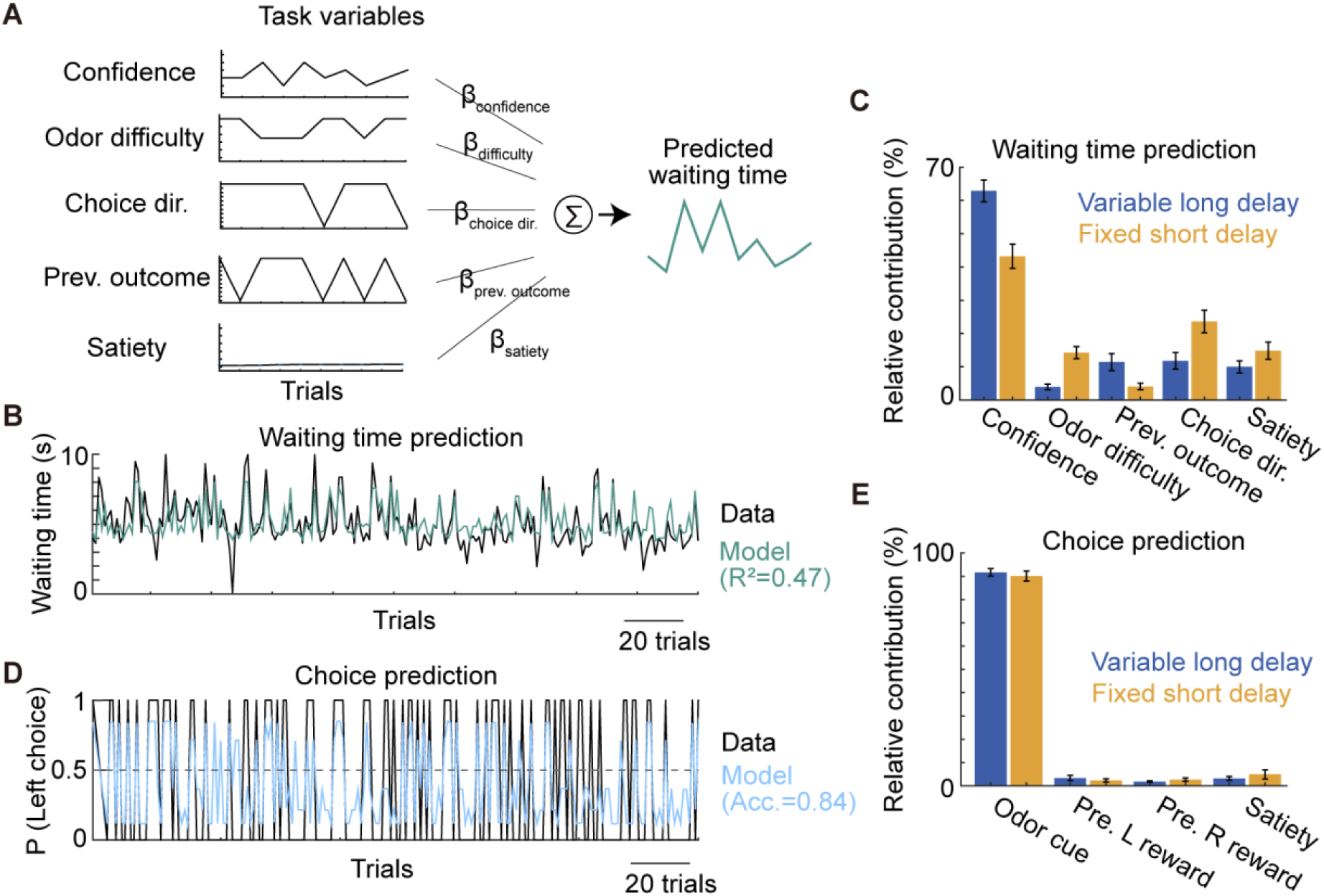
Altered behavioral use of task variables revealed by GLMs. (A) Schematic of the GLM used to predict trial-by-trial reward waiting time based on multiple task factors. (B) Example of GLM-predicted (green) and actual reward waiting time (black) from an example session. (C) Relative contributions of task variables in the GLM predicting reward waiting time. (D) Example of GLM-predicted choice probability (pale blue) and actual choice directions (black) from a single session. (E) Same as C, but for the GLM predicting choice directions. (C & E) Bars show the mean values, and error bars show SEM.

One potential caveat in this interpretation is that the observed differences in waiting time might stem from changes in perceptual decision-making strategies (i.e., odor-guided choices), despite no apparent difference in average choice accuracy (Fig. 1B). To evaluate how perceptual decisions were influenced by various task factors, we employed a logistic GLM to predict the probability of choosing either side based on odor cues, previous reward outcomes for left and right choices (modeled independently), and reward satiety (Fig. 2D). The relative contributions of task variables revealed a predominant influence of odor cues in both session types (Long: 92.6 ± 1.2%; Short: 90.9 ± 1.2%), with minimal contributions from other factors (< 5.6%) (Fig. 2E). These results further suggest that the rats specifically adapted their time-investment strategies without altering their perceptual decision strategies.

### Neurons representing post-decision confidence emerged in OFC regardless of the degree of its behavioral expression

To investigate how changes in behavioral strategy affect confidence representations in the orbitofrontal cortex (OFC), we recorded the activity of hundreds of single neurons during both session types (Long: *n* = 840; Short: *n* = 713). We focused our analyses specifically on the reward waiting time—when rats fix their snout in the chosen port while anticipating the trial outcome—allowing us to directly associate neuronal activity with the time investment decisions. To identify neurons encoding decision confidence, we computed a conventional selectivity index for each neuron using receiver operating characteristic (ROC) analysis. This method quantified the extent to which firing rate distributions differentiated between correct and error trials during the reward waiting period (i.e., outcome prediction selectivity), serving as a proxy for single-neuron encoding of confidence (high vs. low)^32^. Although individual neurons exhibited diverse temporal firing patterns in both session types (Fig. 3A and C), averaging across the neuronal population revealed clear tuning to confidence levels throughout the reward waiting time (Fig. 3B and D). Interestingly, a comparable number of neurons were classified as significantly selective in the short delay sessions relative to the long delay sessions (Fig. 3E and Supplementary Fig. 2C; permutation test, *p* < 0.05). Moreover, the strength of outcome prediction selectivity tended to be higher in the fixed short delay sessions than in the variable long delay sessions (Fig. 3F; Kolmogorov-Smirnov test, *p* = 2.5×10^-5^). This difference in selectivity was not attributable to lower average firing rates in the long delay sessions, which could have been affected by the longer analysis window (Kolmogorov-Smirnov test, *p* = 0.454; Supplementary Fig. 2B for all neurons). Given the reduced behavioral expression of confidence in the short delay sessions (Figs. 1 and 2), the comparable or even enhanced confidence representations in these sessions were surprising.

**Fig. 3:**
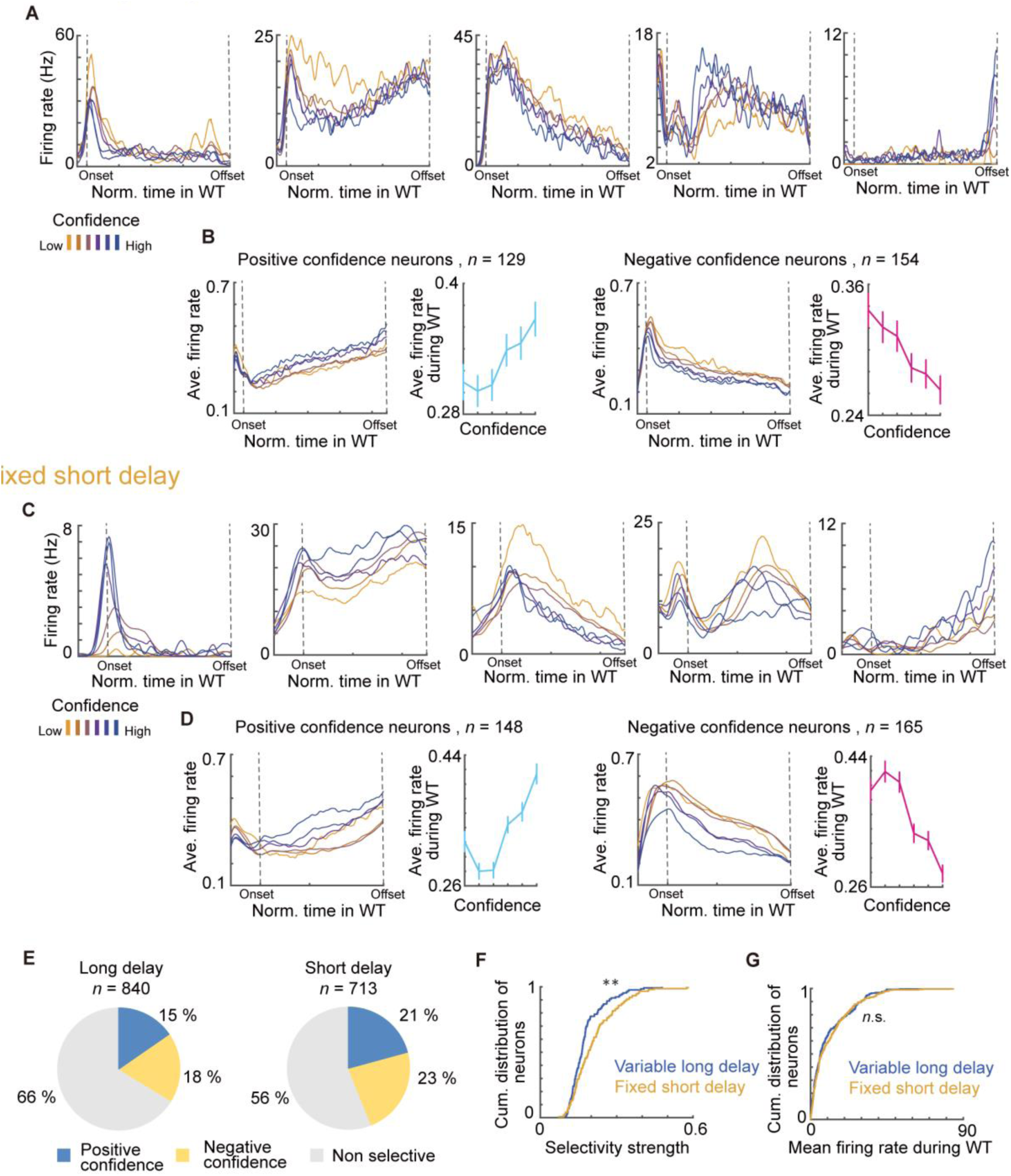
Individual neurons in the OFC represented statistical confidence. (A) PETHs of five example OFC neurons during reward waiting time of the long delay sessions. To facilitate visualization of neuronal activity across trials with different confidence levels, which typically had different durations of trials, time on x-axis is normalized to the same length (Methods). Line colors indicate firing rates averaged across trial types that differ in predicted confidence levels, as estimated by signal detection theory (i.e., based on the interaction between sensory evidence and choice outcome). (B) PETHs and tuning curves of neurons positively (left) and negatively (right) representing confidence levels. The tuning curves are derived from time-averaged firing rates during the waiting time. (C-D) The same as A and B but for the short delay session. (E) Proportion of neurons with significant outcome prediction selectivity during the waiting time in each session type (ROC test with permutation; *p* < 0.05). (F) Comparison of the magnitude of outcome prediction selectivity. Neurons with significant selectivity were included in this analysis. Asterisks show statistical significance (*p* < 0.05; Kolmogorov-Smirnov test). (G) The same as F but for comparison of time-averaged firing rates during the reward waiting time (Kolmogorov–Smirnov test, *p* = 0.454). (B & D) Lines show the mean values, and error bars show the standard error of the mean (SEM).

However, closer examination of peri-event time histograms (PETHs) revealed that the firing rates of neurons with significant selectivity were more precisely aligned with a continuum of confidence levels in the long delay sessions compared to the short delay sessions (Fig. 3B, D). In the short delay sessions, the graded firing patterns were notably reduced at lower confidence levels, resulting in a less linear tuning profile (Fig. 3D) that paralleled the weaker relationship between time investment and confidence observed in Fig. 1G. These findings suggest that although a similar number of neurons were classified as encoding confidence, their tuning precision declined when time investment behavior relied less on confidence (Fig. 2C).

### Behavioral strategies modulated the temporal dynamics of confidence and other task-variable representations

If the activity of OFC neurons is important for guiding time investment decisions, then the temporal dynamics of task-relevant representations in the OFC may also reflect the behavioral strategy being used. Specifically, if confidence representations guide waiting behavior, these signals may be more sustained in the long delay sessions, where reward waiting time is prolonged. Conversely, in the short delay sessions, a timing-based strategy may engage OFC neurons up to the expected reward time, after which task-relevant activity may diminish. To investigate the temporal dynamics of task-variable representations in the OFC, we used a kernel-based GLM (see Methods)^43,44^. This model predicted the instantaneous firing rates of individual neurons based on several task variables. These included decision confidence and two other task variables—odor difficulty and choice direction—which were particularly emphasized in the short delay sessions (Fig. 2C). The kernels captured the timing-dependent effects of each task variable across three distinct behavioral epochs: choice onset (from −0.1 to 0.5 seconds), choice offset (from −0.5 to 0.1 seconds), and the variable interval between these two events (Supplementary Fig. 3A). Additionally, we incorporated a timing-independent kernel for reward satiety, resulting in a total of 10 types of kernels (Fig. 4A).

**Fig. 4:**
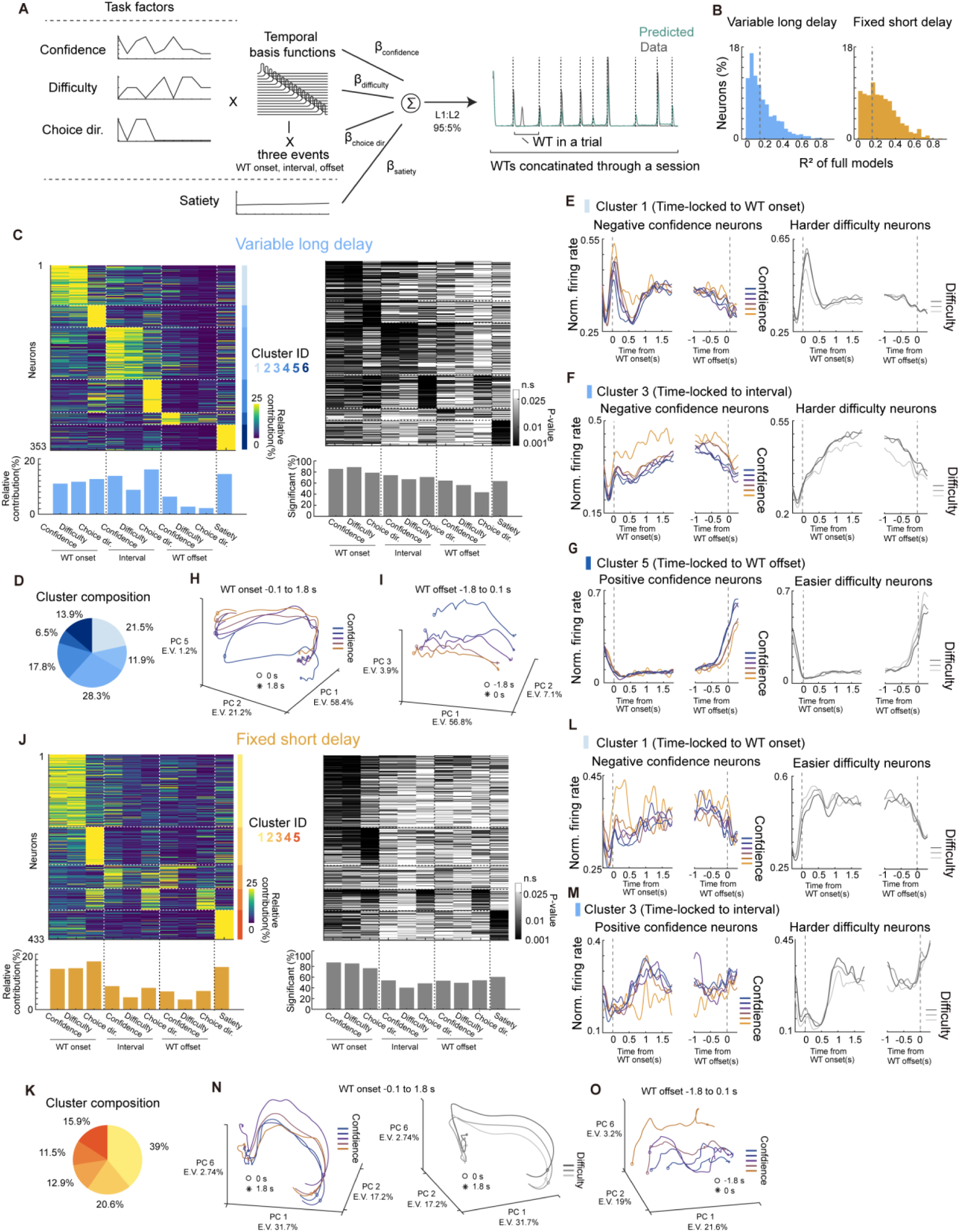
Task encoding dynamics of individual neurons revealed by GLMs. (A) Schematic of the GLM used to predict the instantaneous firing rates of individual neurons, based on kernels representing combinations of task variables and events during the reward waiting time (left). An example of the predicted firing rate of a neuron with its actual activity (green and gray, respectively) is shown on the right. (B) Distribution of the GLM prediction performance in each session type. Gray lines indicate the thresholds used to select neurons for further analysis. (C) The colored heatmap (left) shows clusters of neurons with similar task-related representations, identified by GMM clustering based on relative contributions. Blue bars (bottom left) indicate the average relative contributions of task variables across neurons. The grayscale heatmap (right) shows the significance of each neuron’s selectivity to task variables. The bar plot (bottom right) indicates the percentage of neurons significantly selective to each variable (*p* < 0.025, t-test with FDR correction). (D) Proportion of neuronal clusters in the long delay session. Cluster colors correspond to those shown on the right side of the colored heatmap in C. (E) Average firing rates of neurons negatively tuned to confidence (left) and neurons positively tuned to odor difficulty (right) in Cluster 1, which were temporally locked to the waiting time onset. (F) The same as E, but for Cluster 3, which was temporally locked to the interval during the waiting time. (G) Similar to E and F, but for Cluster 5, which was temporally locked to the waiting time offset. The left panel shows neurons positively tuned to confidence, and the right panel shows neurons negatively tuned to odor difficulty. (H) Trajectory of population activity representing confidence levels aligned to the waiting time onset. (I) The same as H but aligned to the waiting time offset. (J-M) Same as C-F, but for the short delay session. (N) Same as H, but for the short delay sessions. The right panel shows the activity trajectory of the same population as in the left panel but conditioned on the odor difficulty levels. (O) The same as I, but for the short delay sessions.

We grouped neurons with relatively good model fits (*R²* > 0.15; gray lines in Fig. 4B) and functionally similar neurons characterized by relative contributions into distinct clusters using a Gaussian Mixture Model (GMM) for each session type (see Methods)^16,42^. Based on the Bayesian Information Criterion (BIC; Supplementary Fig. 3B), we identified six clusters of neurons in the long delay session, each characterized by specific combinations of task variables and events (top left panel in Fig. 4C). Neurons encoding confidence and odor difficulty (Clusters 1, 3, and 5) tended to form separate clusters from those encoding choice direction and reward satiety (Clusters 2, 4, and 6, respectively), suggesting the presence of distinct clusters for different task variables. Furthermore, most clusters were temporally aligned with either the waiting time (WT) onset (Clusters 1 and 2; 33.4% in total) or the WT interval (Clusters 3 and 4; 46.1%), while fewer clusters were aligned with WT offset (Cluster 5; 6.5%) (Fig. 4C– D). Interestingly, neurons that encoded both confidence and odor difficulty tended to exhibit more clearly graded tuning to confidence levels than to odor difficulty levels (Fig. 4E–G). We recapitulated the activity of all the clusters as a population using principal component analysis (PCA; see Methods). Following WT onset, population activity was largely dominated by temporal dynamics (PC1 and PC2 on the x- and y-axes in Fig. 4H). Although it accounted for a relatively small portion of the explained variance (1.2%), we identified a dimension that captured confidence levels in a temporally stable manner (PC5 on the z-axis). Notably, this confidence-related dimension explained a greater proportion of variance (PC2: 7.1%, PC3: 3.9%) when neuronal activity was aligned to WT offset (Fig. 4I), suggesting a gradual increase in the dominance of this signal throughout the interval period. These findings suggest that, when confidence guides time investment decisions, it is sustained over time through the recruitment of neurons with distinct temporal response profiles (Fig. 4C).

The composition of neuron clusters in the short delay sessions exhibited distinct patterns compared to the long delay sessions. Although the number of clusters (*n* = 5) and the separation between confidence-/odor-difficulty–related, and choice-direction–related clusters remained consistent (Fig. 4J), most neurons were temporally aligned with the onset of the WT, with Clusters 1 and 2 together comprising 59.6% of the population (Fig. 4K). Since the WT onset kernels capture neuronal activity during the first 0.5 seconds of WT, the dominance of these clusters suggests that the activity of many neurons is time-locked to the expected reward timing at 0.5 seconds. In contrast, neurons associated with the interval period (Clusters 3 and 4) accounted for only 24.4%, nearly half the proportion observed in the long delay sessions. Interestingly, neurons encoding confidence and odor difficulty exhibited less distinct graded tuning to confidence levels, particularly during the interval period (left panels in Fig. 4L–M), but showed clearer tuning to odor difficulty (right panels in Fig. 4L–M). Although population activity differentiated between confidence levels after WT onset, the trajectory was not aligned along a gradient from low to high confidence (left panel in Fig. 4N). Instead, it more closely followed odor difficulty levels (right panel in Fig. 4N). Notably, confidence levels were more clearly represented in a graded manner when neuronal activity was aligned to the WT offset, despite temporal variability in the trajectory (PC6 on the z-axis in Fig. 4O). This suggests less precise and temporally unstable confidence representations in the short delay sessions, contrasting with our findings of stable and precise confidence representations in the long delay sessions (Fig. 4H-I). Taken together, the GLM analysis revealed that the recruitment of OFC neurons is strongly dependent on the behavioral expression of confidence, which is shaped by task demands.

### The recruitment of confidence-related neuronal clusters reflected the extent to which rats relied on confidence to guide their behavior

We next aimed to examine how the individual functional neuronal clusters revealed by the GLM, relate to the behavioral expression of confidence. To examine this relationship, we computed the correlation between the average relative contributions of each variable across simultaneously recorded neurons and the M-ratio (*d’*/meta-*d’*, as defined in Fig. 1K) for each session. In the long delay sessions, we observed relatively strong positive correlations between the contributions of confidence (*r* = 0.46, *p* = 0.0002) and odor difficulty (*r* = 0.48, *p* = 0.0001) during the interval period and the M-ratio (Fig. 5A). Interestingly, this positive correlation between neuronal activity and metacognitive efficacy was specific to that particular combination of task variable and event timing. Neither confidence nor odor difficulty showed similar correlations during the WT onset or offset periods, nor did choice directions during the interval period (Fig. 5B). We then performed a similar analysis to examine the relationship between the cluster composition and M-ratio in each session. Strikingly, Cluster 3—which primarily represented confidence during the interval period (Fig. 4C)—showed a positive correlation with M-ratio (*r* = 0.33, *p* = 0.0085; Fig. 5C-D). In contrast, the other clusters showed either lower or negative correlations, highlighting that greater recruitment of neurons representing confidence during the extended interval in WT is associated with more efficient behavioral expression of confidence. We performed the same analysis for neuronal activity in the short delay sessions. Interestingly, we observed a similar trend: only confidence coding during the interval period, along with the proportion of the cluster corresponding to this activity (Cluster 3), showed positive correlations with the M-ratio (Fig. 5E-H). Given that the relative contribution of confidence and the proportion of this cluster were higher in the long delay session (Fig. 4C & J), these results further suggest that sustained confidence representation during WT is crucial for the efficient behavioral expression of confidence.

**Fig. 5:**
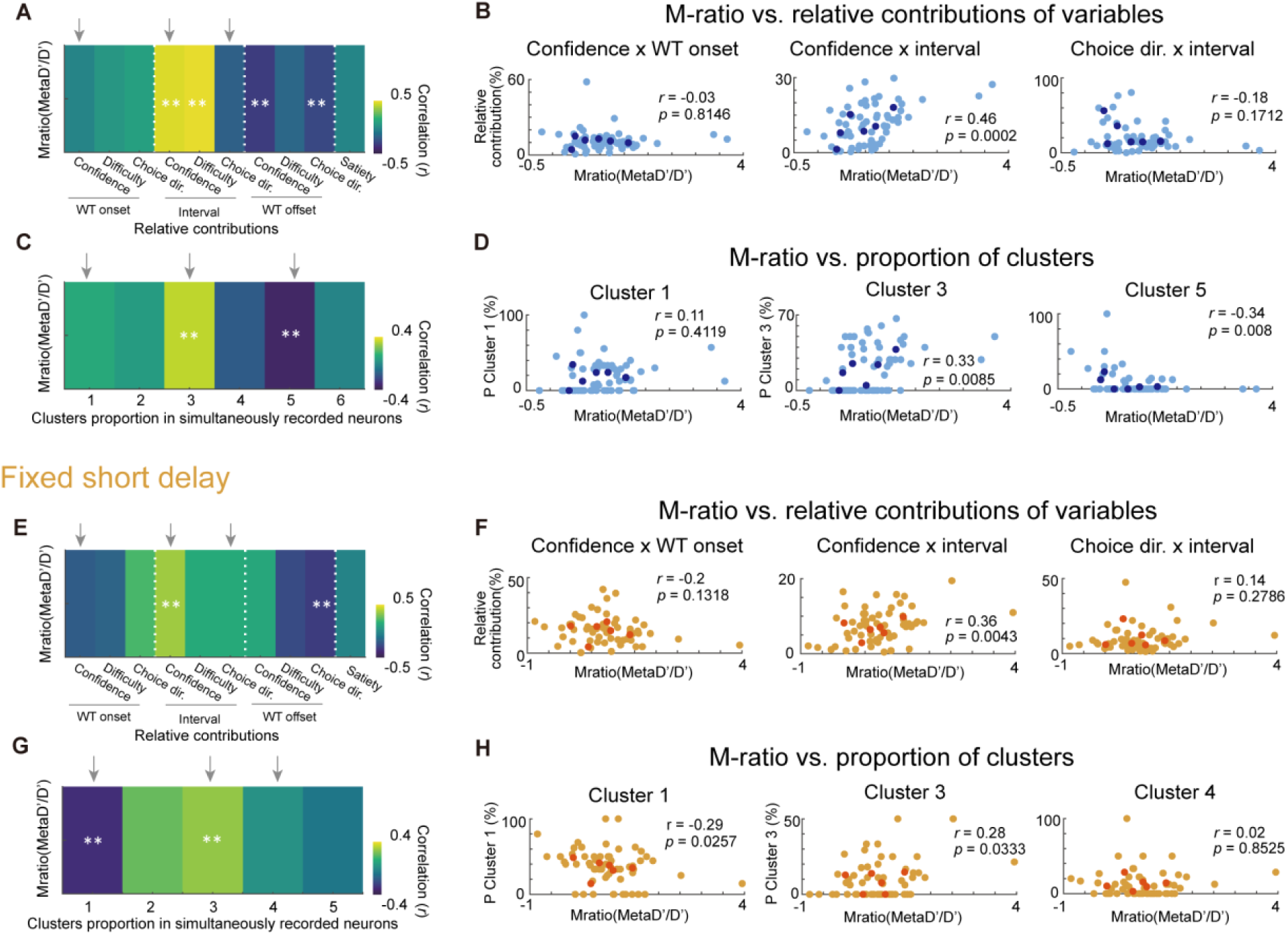
Relationship between task encoding and efficient behavioral expression of confidence. (A) Heatmap showing the correlation between the average relative contributions of task variables across neurons and the M-ratio across long delay sessions. Gray arrows indicate the conditions shown further in detail in B. (B) Scatter plots showing the relationship between the average relative contribution of each task variable and the M-ratio across sessions, corresponding to the data underlying the correlations shown in A. Pale dots represent individual sessions, and darker dots indicate session averages for each animal. Correlation values were computed across sessions. (C) The same as A but showing the correlation between the proportion of each neuronal cluster and the M-ratio across long delay sessions. (D) The same as B, but the relationship between each cluster’s proportion and the M-ratio across sessions. (E-H) The same as A-D, but for short delay sessions. In heat maps, white asterisks indicate significant correlations (*p* < 0.05).

### Behavioral expression of decision confidence is associated with temporally stable population decoding

So far, we have demonstrated that confidence coding—particularly during the extended duration of the WT—is associated with the behavioral expression of confidence. While PCA indicated more temporally stable confidence representations at the population level when confidence was more strongly expressed behaviorally (i.e., in the long delay compared to the short delay sessions), we sought to quantify this coding stability and examine how the different clusters contributed to it. To this end, we performed a time-resolved decoding of confidence from OFC population activity (*n* = 334 neurons) using a linear support vector machine (SVM; see Methods). We trained a binary decoder to classify relatively high and low confidence levels (i.e., correct vs. error trials) based on population activity at each time point during the WT.

As shown in Fig. 6A, binary confidence levels were successfully readout from OFC population activity in the long delay sessions. Decoding performance rose around the WT onset and remained relatively stable at a high level. After WT offset, decoding accuracy stayed above chance briefly but dropped to chance levels within 0.5 seconds. To examine how the underlying population encoding patterns evolved over time, we also performed cross-temporal decoding^45,46^. A transient encoding state emerged just before WT onset and persisted for about 0.5 seconds afterward (left panels in Fig. 6B-C). This was followed by a temporally generalized encoding state, indicating stable confidence representations during the middle of the WT period. This stable state was also evident when activity was aligned to WT offset but sharply diminished just before the offset. We applied the same analysis to population activity in the short delay session. While decoding accuracy remained above chance during the WT, it exhibited greater temporal variability and declined to chance levels even before the WT offset (Fig. 6D). Cross-temporal decoding revealed an initial transient encoding state, similar to that observed in the long delay sessions; however, the subsequent pattern was less temporally generalized (Fig. 6E-F), indicating reduced stability of confidence representation during the WT.

**Fig. 6:**
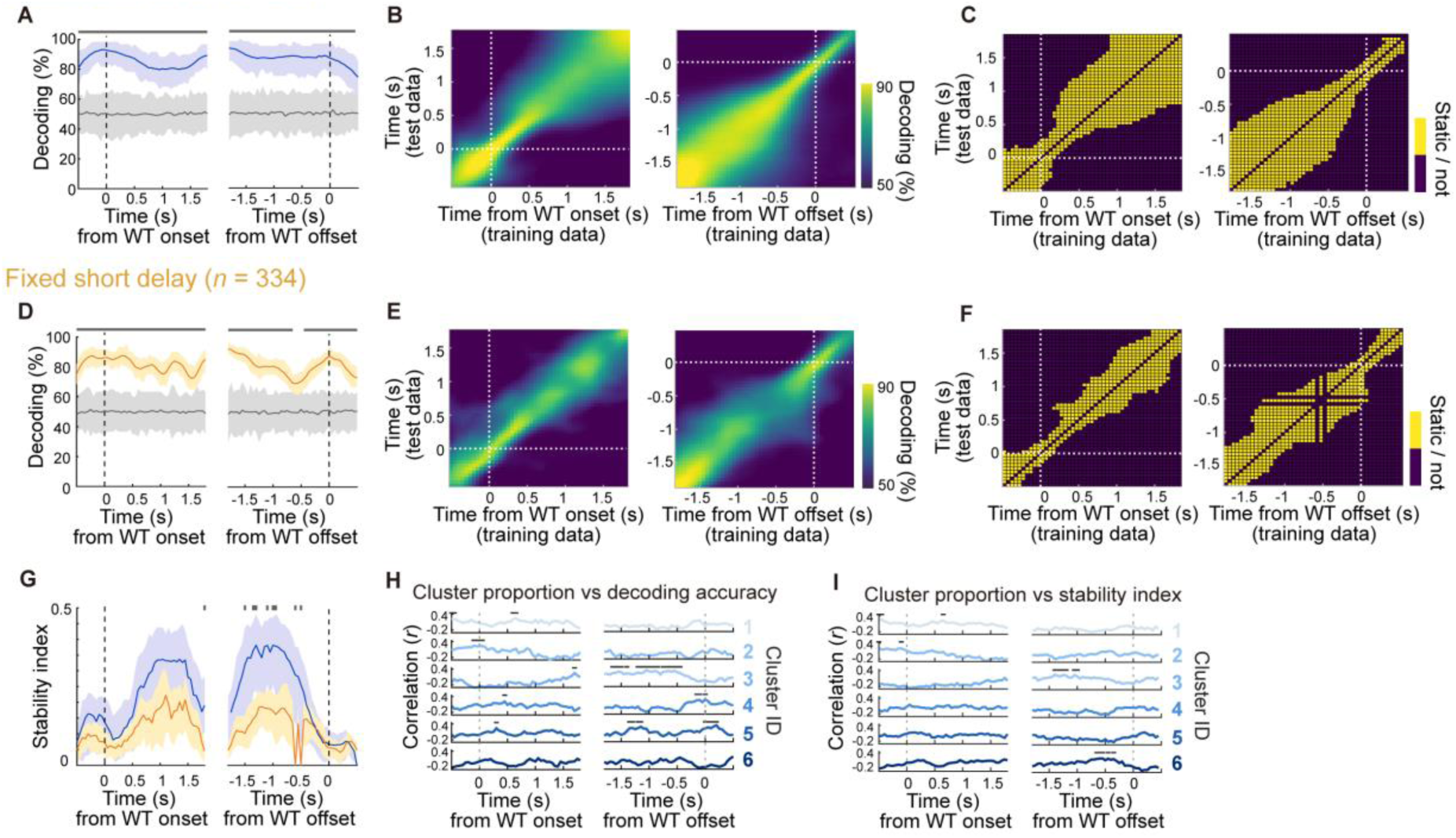
Time-resolved decoding of confidence during the reward waiting time from the OFC population activity. (A) Time-resolved population confidence decoding performance in the long delay sessions. Solid lines indicate the average decoder performance, and shaded areas represent the 95% percentile range obtained from iterations (blue: real data; gray: label-shuffled). The gray bar above the plot marks time points with significantly above-chance decoding performance (estimated *p* < 0.05; see Methods). (B) Cross-temporal decoding performance corresponding to the decoder shown in A. On-diagonal data points represent decoding accuracy when training and testing on the same time bins (also shown in A). Off-diagonal data points represent decoder performance when testing on time bins different from the training bins, indicating the temporal generalization of the decoder. Decoding performance around the waiting time onset (left) and offset (right) are shown independently. (C) Stability of the cross-temporal decoding performance shown in B. Yellow marks time bins where the decoder performance does not significantly decrease from the on-diagonal performance, indicating temporally stable representations. (D-F) Same as A-C but for the short delay session. (G) Comparison of the temporal stability of decoding performance between the long and short delay sessions. The gray bar above the plot indicates time points with significant difference (estimated *p* < 0.05; see Methods). (H) Correlation between decoding performance (on-diagonal values from cross-temporal decoding) and the proportion of GLM-defined neuronal clusters from Fig. 4 in the long delay sessions. The gray horizontal bar marks time points with a significant positive correlation (*p* < 0.05). (I) the same for G but for correlation between the proportion of GLM-defined neuronal clusters and stability index of decoding performance.

To quantify the temporal generalization of coding patterns, we computed a stability index at each time point^45,47^. This index increased during the middle of the WT and was significantly higher in the long delay sessions than in the short delay sessions (Fig. 6G). Notably, the higher stability index was especially prominent when neuronal activity was aligned to the WT offset, suggesting that stable coding gradually emerged over the course of extended WT in the long delay sessions. To further examine the basis of this stable coding, we leveraged the subsampling of neurons from the long delay sessions used in each decoding iteration. We analyzed the contributions of different functional clusters (Fig. 4D) to confidence decoding. Interestingly, the proportion of neurons from specific clusters included in each decoding iteration positively correlated with decoding accuracy and the stability index in distinct ways (black bars in Fig. 6H-I). In particular, higher proportions of Cluster 3 neurons were associated with greater decoding accuracy and higher stability index during the middle of WT. Other clusters also showed positive correlations, typically aligned with the temporal epochs in which they were most active (e.g., Cluster 1 just after WT onset). Taken together, these results indicate that functional clusters of neurons collectively form temporally stable representations of confidence.

### Behavioral expression of decision confidence is associated with linearized population coding of confidence

The remaining question is how graded levels of confidence are represented at the population level, and how the geometry of these representations relates to behavioral expression of confidence. To investigate this, we visualized the representational geometry encoded by OFC neuronal populations by performing PCA on population activity across task conditions defining confidence levels (odor types and choice outcomes), time-averaged during the WT^28,48,49^. In the long delay sessions, these conditions formed an X-shaped structure in the reduced space (Fig. 7A), consistent with a continuum of confidence levels. This structure was well captured along a single linear dimension (PC2; Fig. 7B). Given that PCA is an unsupervised method, the emergence of a dedicated linear axis representing confidence was striking. Additional principal components (PCs) corresponded to distinct task variables—PC1, PC3, and PC4 encoded choice, odor identity, and difficulty, respectively—indicating that the OFC encodes multiple task variables in a disentangled (or factorized) fashion ^23,27,50^. In contrast, the representational geometry observed during the short delay sessions did not exhibit such an organized structure (Fig. 7C). Notably, PC2 in the short delay sessions (Fig. 7D) resembled the U-shaped pattern of reward waiting time observed behaviorally (Fig. 1H), consistent with the reduced expression of confidence in these sessions (Fig. 1I–K). Moreover, PCs 3–4 in the short delay sessions showed weaker alignment with task variables, suggesting that the degree of representational disentangling in the OFC is influenced by behavioral strategy.

**Fig. 7:**
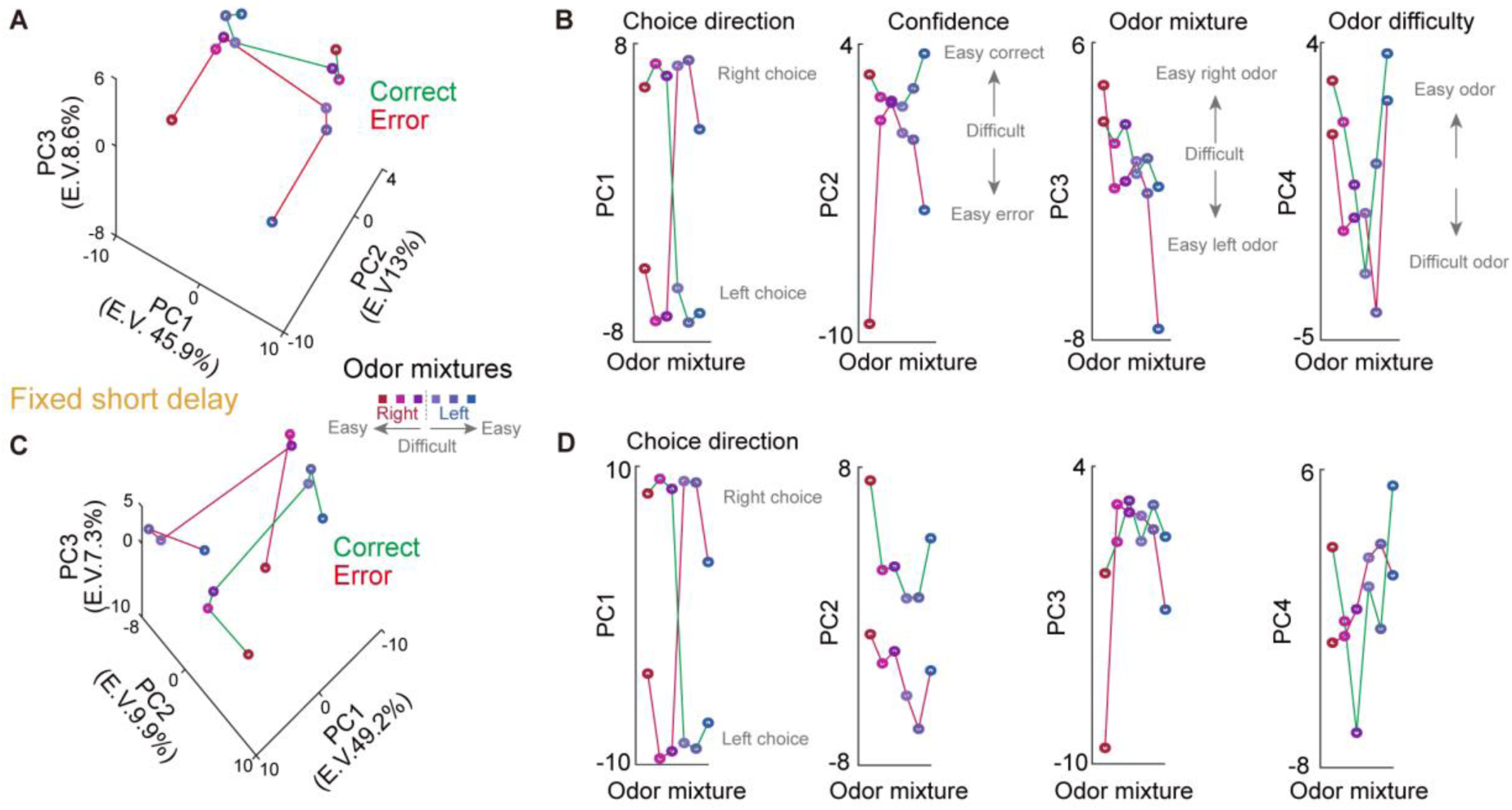
Distinct geometry of task-variable representations induced by long and short delay sessions. (A) Three-dimensional PCA plot of population activity across task conditions relevant to confidence. Circle and line colors represent odor type (six odor mixtures) and choice outcome (correct or error), respectively. (B) Population activity projected onto each PC axis. Each axis captures intuitively interpretable task variables such as choice direction (PC 1), confidence (PC 2), odor mixtures (PC 3), and odor difficulty (PC 4). (C-D) The same as A-B, but for the short delay session. Each PC axis captures patterns that are less interpretable in terms of task variables.

Since a key advantage of disentangled representations is their linear accessibility for downstream readout leading to behavioral outputs, we directly tested whether a continuum of confidence levels could be linearly decoded using a support vector machine (SVM) (see Methods). We trained the decoder to classify the two lowest levels of confidence (easy + medium odor error vs. difficult odor error trials) and then tested its ability to classify both the lowest and higher levels (i.e., correct trials) of confidence (generalization; Fig. 8A). This approach predicted that the decoder’s generalization performance should be better when the confidence representation is more linearized and would drop when the representation exhibits curvature, potentially induced when the confidence representation is tangled with relevant another variable such as odor difficulty levels (Fig. 8B-C). In the long delay session, decoder performance gradually increased toward higher confidence levels in the generalized conditions (Fig. 8D), consistent with predictions from a linearized representation of confidence. In contrast, the short delay session exhibited a different pattern. Although the decoder reliably classified the lower confidence levels it was trained on, its performance sharply decreased as the confidence level increased in the generalization conditions, reaching the chance level (Fig. 8E). This poor decoder performance was not due to a lack of population activity differentiating confidence levels, as one-by-one decoding showed generally higher decoding performance in the short delay session than in the long delay session (Fig. 8F).

**Fig. 8:**
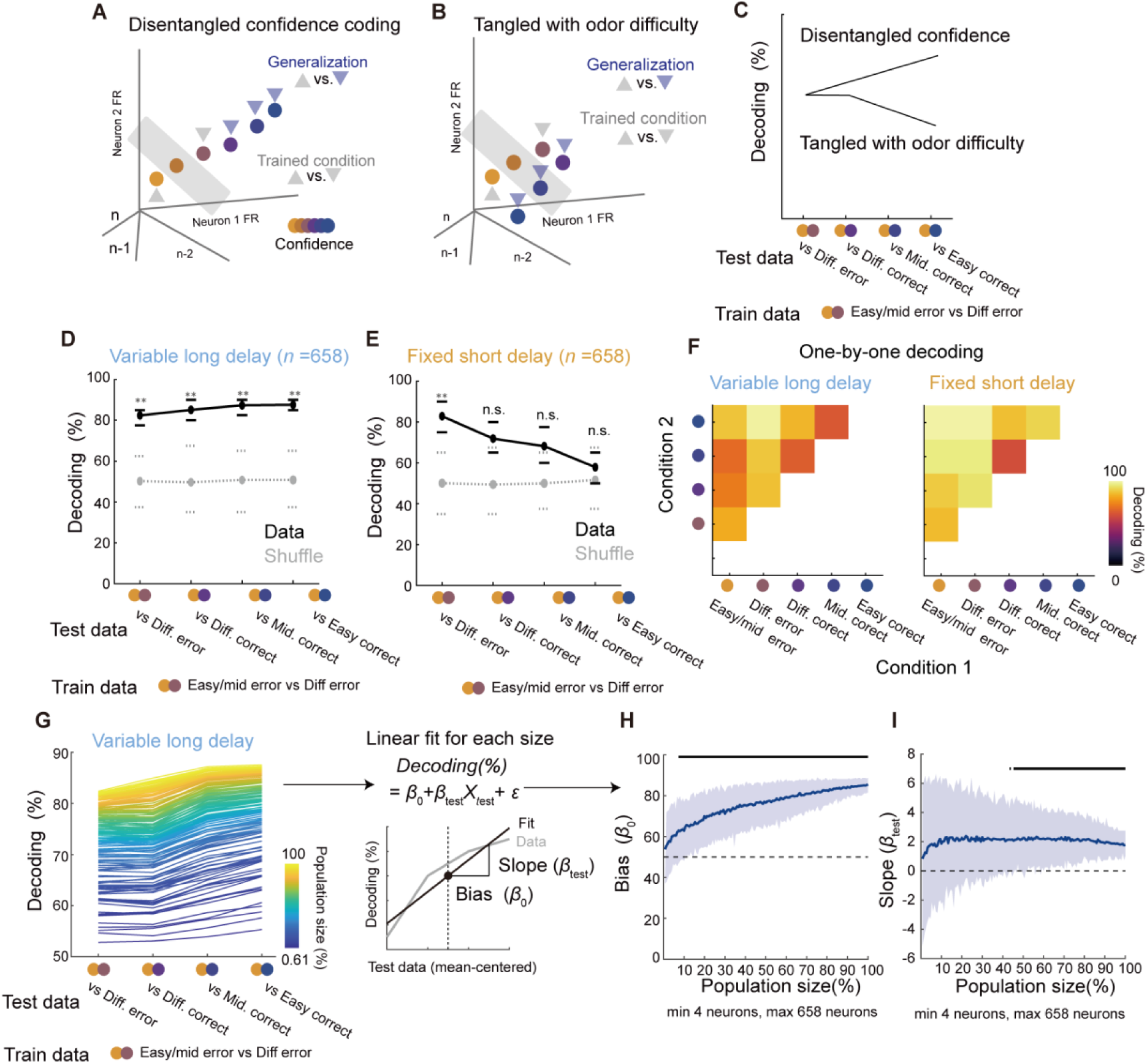
Linearization of confidence representation revealed by population decoder generalization. (A) Schematic of a linearized confidence representation that enables generalization of a linear decoder. A decoder trained on low-confidence trials (two gray triangles) establishes a hyperplane (gray rectangle) that successfully classifies between the lowest confidence level (upward gray triangle) and higher levels (blue triangles). (B) Schematic of a non-linear confidence representation, where curvature in the geometry—reflecting confidence tangled with odor difficulty—impairs the generalization capability of a linear decoder. (C) Predicted generalization performance of decoders on linear versus non-linear confidence representations, as illustrated in A and B. (D) Decoder generalization performance in the long delay session. Circles indicate average performance, and lines show the 95% range across iterations (black: real data; gray: label-shuffled control). Asterisks indicate above chance decoding performance (estimated *p* < 0.05; see Methods). (E) Same as D, but for the short delay session. (F) One-vs-one decoding performance averaged across iterations for the long (left) and short (right) delay sessions. (G) Decoder performance across different population sizes, averaged across iterations. The right panel illustrates the method for independently evaluating overall decoding performance and decoder generalization using a linear fit of the obtained decoder performances. (H) Relationship between overall decoding performance (*β_0_*) and population size. Black bars indicate statistically significant performance compared to 50% chance level (estimated *p* < 0.05). (I) Same as H, but for decoder generalization. Black bars indicate statistically significant generalization (*β_test_* > 0), evaluated using estimated *p*-values. (H-I) Lines indicate mean decoder performance, and error bars represent SEM.

We also tested the importance of population size for decoding accuracy and generalization by analyzing neuronal populations of varying sizes (*n* = 4 to 658 neurons) from the long delay sessions. We quantified both overall decoding performance and generalization performance for each population size using the bias (*β₀*) and slope (*β_test_*) of a linear fit (Fig. 8G). General decoding performance increased monotonically with population size and exceeded chance levels with relatively few neurons (7.3%, 48 neurons; Fig. 8H). Interestingly, the minimum number of neurons required to match the performance of the full population was approximately 20% (132 neurons), suggesting that confidence can be reliably decoded from a moderate-sized population of OFC neurons. Generalization performance also plateaued quickly with a small number of neurons, although increasing the population size further reduced variability across iterations (Fig. 8I), indicating that reliable generalization can be achieved by a subset of neurons. Taken together, these findings demonstrate that confidence can be robustly and linearly decoded from a moderately sized OFC population, particularly when it is strongly reflected in behavior.

Finally, we examined whether confidence decoding could predict differences in the behavioral expression of confidence across sessions. For each session with 5 or more neurons, we used simultaneously recorded neuronal populations to quantify both decoding accuracy and generalization performance. We observed clearer generalization slopes in the long delay sessions (Fig. 9A). Although the overall decoding performance—as indicated by the bias (*β₀*) of the linear fit—was comparable across session types (Fig. 9B; t-test, *p* = 0.191), the slope (*β__test_*), which reflects generalization, was significantly higher in the long delay sessions (Fig. 9C; t-test, *p* = 0.001). Furthermore, the long delay sessions with relatively high decoding accuracy (*β₀*) exhibited higher M-ratios than those with lower decoding accuracy (Fig. 9D; t-test, *p* = 0.016), indicating that general confidence decoding accuracy predicts more efficient behavioral expression of confidence. This relationship was not observed in the short delay sessions, suggesting a decoupling between confidence representation and its behavioral expression when task demands are lower (t-test, *p* = 0.398). Similarly, in the long delay sessions, positive decoder slopes—reflecting better generalization—were associated with higher M-ratios than negative slopes (Fig. 9D; t-test, *p* = 0.007). Again, this association was absent in the short delay sessions (t-test, *p* = 0.085). These findings suggest that the extent to which confidence representations can be linearly accessed determines their impact on behavior, especially when task demands are high.

**Fig. 9:**
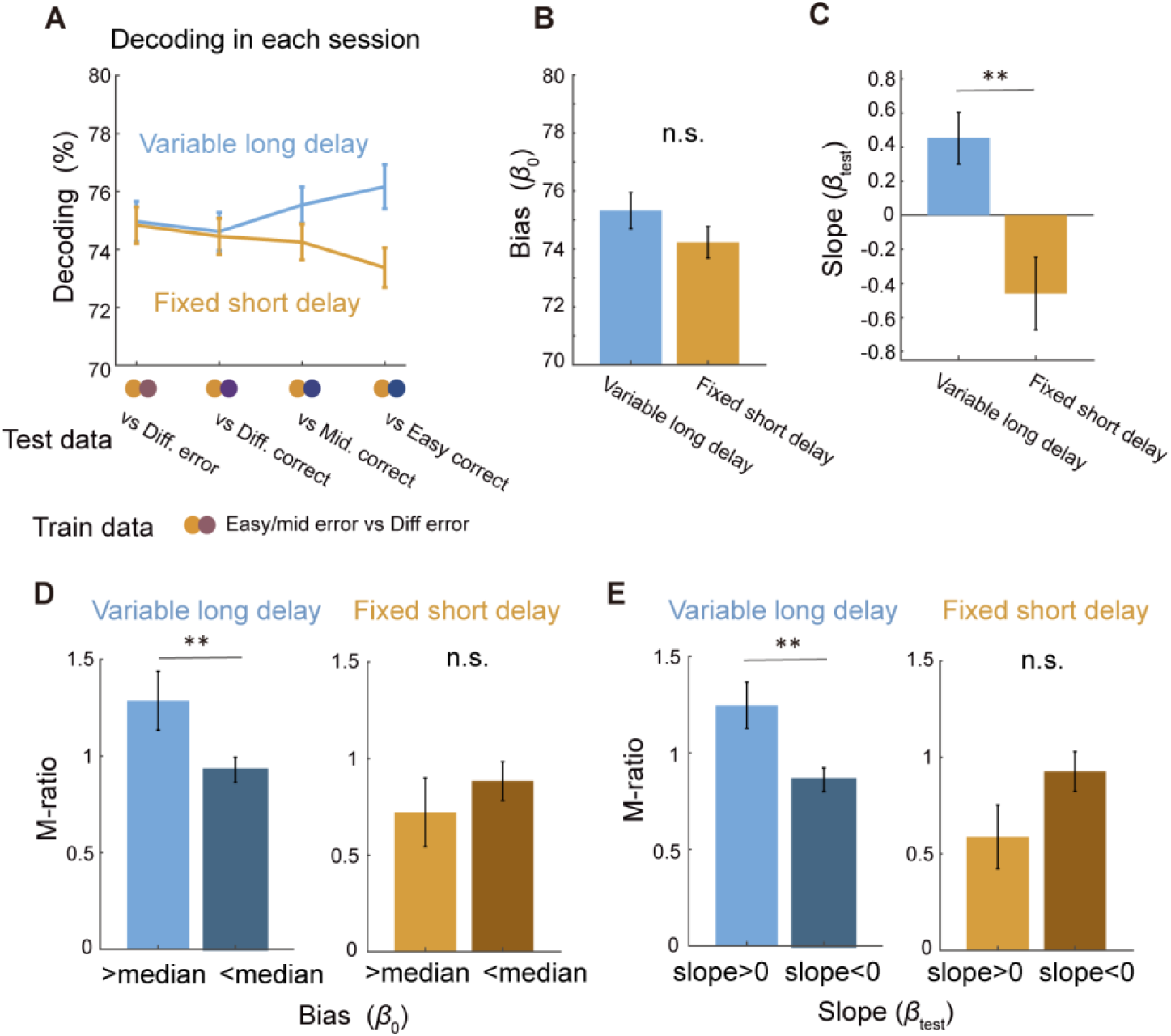
Confidence decoding performance predicts the efficient behavioral expression of confidence. (B) Decoder generalization performance using simultaneously recorded neurons from each recording session. (B–C) Comparison of the decoder bias (*β₀*) and slope (*β_test_*) obtained from simultaneously recorded neurons between session types. (D) Comparison of metacognitive performance (M-ratio) between sessions with higher and lower general decoding performance (*β₀*), defined relative to the median β₀ within each session type. (E) Same as D but comparing M-ratio between sessions with higher versus lower decoder generalization, as defined by the decoder slope (*β_test_*). (B–E) Colored bars indicate session average, and error bars represent SEM. Asterisks indicate statistical significance (t-test, *p* < 0.05); *n.s.* indicates non-significance.

## Discussion

Our findings demonstrate that confidence coding in the OFC tracks the behavioral expression of confidence. Interestingly, the proportion of confidence-encoding neurons remained stable, even when the behavioral expression of confidence varied with task demands (Fig. 1–3). However, functional clustering revealed a reorganization of the temporal dynamics of these neurons (Fig. 4 & 5), and confidence representations became more temporally stable at the population level when confidence was strongly expressed in behavior (Fig. 6). Additionally, the linear geometry of population representations across the confidence spectrum was tightly coupled with its behavioral expression (Fig. 7–9).

A conventional approach to demonstrating the involvement and functional significance of a given brain region in cognitive functions has been the identification of neurons whose activity reflects task-relevant information. Based on this view, it has been predicted that behavioral engagement of a specific cognitive process would recruit neurons to represent relevant information. This framework is supported by numerous studies, particularly those focusing on task acquisition and task engagement^1–14^. From this perspective, our finding of a comparable proportion of confidence neurons—regardless of their behavioral expression—was unexpected (Fig. 3). A key strength of our study was the consistent task design, which allowed us to control the rats’ behavior (i.e., by requiring them to fix their snout in the chosen port during anticipating reward), while still inducing different levels of behavioral use of confidence between long and short delay sessions (Fig. 1 & 2). This design, combined with an encoding model of neuronal activity, revealed temporal dynamics that reflected distinct behavioral strategies: confidence- vs. timing-based decisions (Fig. 4). These results suggest that differential engagement of neurons in the OFC may flexibly support distinct computations depending on demands from contexts. Although previous studies implicating the OFC in both functions support this interpretation^29–32,51–53^, it remains unclear how the OFC supports such flexibility on shorter timescales such as minutes depending on evolving context. Additionally, the OFC is implicated in a variety of functions beyond these two, such as motivation^54,55^, devaluation^52,56^, risk assessment^57^, and reversal learning^58,59^. A significant open question is how the OFC flexibly shifts between these diverse roles despite its limited neuronal resources.

One potential solution to cognitive multitasking, extensively discussed from a computational perspective, is the use of disentangled (or factorized) representations, which enable the linear readout of multiple cognitive variables simultaneously through population activity^23,27,50,60^. These representations have been widely studied as a hallmark of higher-order cognitive systems in both the brain and artificial neural networks^25^. While recent studies in humans and non-human primates have highlighted the importance of such population representations for generalizing cognitive performance across behavioral contexts^23,27,61^, the representational capacity of rodent cortical populations in this framework remains largely unexplored. Our findings suggest that the orbitofrontal cortex (OFC) in rats exhibits disentangled representations not only for externally defined variables—such as odor cues and choice directions—but also for internally generated cognitive variables, including confidence and odor difficulty. Previous studies have reported that OFC neurons form functional clusters, each preferentially encoding specific decision variables such as confidence or chosen value^31,62^. While these earlier findings did not directly test factorized representations, they point to a potential organizational structure that could support such representations. Our results extend these insights by showing that these neurons representing decision variables exhibit distinct temporal dynamics depending on task demands, and form disentangled representations at the population level.

The degree of disentangling is expected to key in flexibly realizing multiple functions depending on contexts with limited numbers of neurons. We typically recorded from the same depth in the OFC across consecutive sessions with different strategies to facilitate comparison (Method; Table 1); however, whether and how the same neurons shift their activity depending on behavioral strategy and how disentangled representations emerge remain an important question for future studies, particularly those using approaches that track individual neurons across days^63–65^.

From both computational and neuroscience perspectives, linearly decodable population representations within a brain region are thought to facilitate the transmission of information in a form that can be readily interpreted by downstream regions through the summation of synaptic inputs^66,67^. While this suggests that the behavioral readout of cognitive variables could rely on similar mechanisms, it has remained unclear whether task demands that require behavioral output actively promote the linearization of cognitive representations. Our results demonstrated enhanced linearity in confidence representations when they were behaviorally expressed, supporting this possibility. Such representations could potentially be read out by downstream regions more directly involved in action selection, such as the striatum^31,68,69^. Future studies could address this possibility by manipulating specific output pathways of the OFC using optogenetic and chemogenetic approaches.

## Method

### Subjects

Seven male Long-Evans rats (Shimizu Laboratory Supplies, Kyoto, Japan) were individually housed and maintained on a laboratory light/dark cycle (lights on 8:00 A.M. to 9:00 P.M.). Rats were placed on water restriction with ad libitum access to food. The animals were maintained at 85% of their baseline weight throughout the experiments. All experiments were implemented in accordance with the guidelines for the care and use of laboratory animals provided by the Animal Research Committee of the Doshisha University with its approval.

### Behavioral apparatus

The behavioral apparatus (Fig. 1A) has been previously described^28,31,43^. An operant chamber (O’Hara, Tokyo, Japan) with three ports in the front wall for nose-poke responses was enclosed in a soundproof box (Brain Science Idea, Osaka, Japan). Each port was equipped with an infrared sensor to detect the animals’ nose-poke responses. Cue odors were presented via the central port through a stainless tube. The odors were mixed with pure air to produce a 1:10 dilution at a flow rate of 1L/min using a custom-built olfactometer (AALBORG, Orangeburg, NY). Water rewards were delivered from gravity-fed reservoirs regulated by solenoid valves (The Lee Company, Westbrook, CT) through stainless tubes placed inside of the left and right ports. We controlled stimulus presentation and reward deliveries and measured behavioral responses using Bpod and Pulse Pal (Sanworks, Stony Brook, NY).

### Decision-making task with distinct reward delay distributions

The current study modified a behavioral task previously designed to measure post-decision confidence^30,32^. Each trial began when the rats poked their snout into the central port in the operant chamber. After a variable delay (200–600 ms, drawn from a uniform distribution), a binary mixture of two pure odorants (S(+)- and R(−)-2-octanol) was presented, randomly chosen from six different concentration ratios. If the rats successfully kept their nose in the central port for 0.4 s after the cue onset, they were allowed to withdraw and choose either the left or right port. The withdrawal earlier than 0.4s resulted in the cancellation of the trial and starting the next trial. Only choices according to the dominant odorant in the mixture were rewarded with a drop of water (25 µl) after a delay period. Any feedback was given when rats choose the wrong side (i.e. error choice).

We manipulated the length and uncertainty of the reward delivery timing by implementing two session types with distinct reward delay distributions. Each rat experienced both types of reward delay distributions across dairy sessions (Fig. 1C). In the variable long delay sessions, the reward delay was drawn from an exponential distribution with a time constant of 1.5 s, shifted by 0.8 s, and truncated at 8 s. Previous studies have shown that exponential delay distributions produce a relatively constant hazard rate^30,32,35^. In our task, the variable long delay introduced uncertainty in reward delivery timing and increased the cost of waiting, prompting rats to decide whether to wait or forgo the potential reward in each trial based on their post-decision confidence (i.e., time investment decision)^30,32^. In contrast, the fixed short delay sessions used a reward delay of 0.5 s, allowing rats to predict the timing of reward delivery. Previous studies have demonstrated the scalar timing law, which states that interval timing estimation error increases with the duration of the interval. Guided by this principle, we selected a sub second fixed delay to facilitate optimal reward timing anticipation^37^, which was expected to reduce the utility of post-decision confidence in time investment decisions. During the recording sessions, rats were typically alternated between sessions with the variable long delay and the fixed short delay across days (Table 1). Note that one rat performed the task in a block design, completing several sessions under the short delay condition followed by sessions under the long delay condition.

### Training procedure

Rats underwent the step-by-step training described below before surgery for the electrophysiological recordings. The overall training process typically took 1-2 months to complete.

1. Rats were on water restriction to sufficiently motivate them to collect reward in the task.
2. Next, the rats were trained to poke their snout into the central port and then immediately choose either the left or right port to obtain a 20 µl water reward. At this stage of training, they received a reward regardless of which port they chose after the central port poke. This step typically took a day.
3. Next, the rats were required to keep their snout fixed in the central port before withdrawing and choosing either side port. The duration of the central port fixation was gradually extended to 1.2 seconds across trials. This step typically took several days.
4. After confirming that the rats had learned to fix their snout in the central port before responding to either side port, we introduced odor cues (S(+)- and R(−)-2-octanol). Initially, we presented only easy odor mixture cues (95:5/5:95 mixtures). The rats were required to poke their noses into the central port, followed by a cue delay and odor presentation. During both the cue delay (200–600 ms, variable) and the cue presentation (>400 ms), the rats had to keep their snout fixed in the central port to ensure that they received the odor cue. Rats were then allowed to choose either the left or right port by withdrawal from the central port. Only correct choices, according to the presented odor cue, were immediately rewarded by a drop of water. To facilitate task learning, we introduced correction trials, which repeated the same trial until the rats chose the correct side port.
5. After confirming that rats discriminated the easy odor cues well, we introduced reward delay. The reward delay was gradually extended across trials and sessions. To avoid unwanted bias from training history, we grouped rats into two groups for learning reward delay. The first group (*n* = 4 rats) was trained with the variable long delay used in the task. In this group, the fixed reward delay was gradually extended from 0 to 0.8s. Subsequently, the maximum range of the exponentially distributed variable delay gradually increased to 8s, while maintaining a minimum range of 0.8s. The second group (*n* = 3 rats) was trained with the fixed short delay used in the task. The fixed reward delay was gradually extended from 0 to 0.5s during training.
6. In addition to the two easy odor cues, two of each medium and difficult odor mixtures were introduced for medium and difficult cue trials. To keep the rat engaged in the task, we initially used odor mixtures with sufficiently strong contrasts for both medium (e.g., 80:20/20:80 mixtures) and difficult cues (e.g., 70:30/30:70 mixtures). The odor contrasts for medium and difficult cues were gradually adjusted to maximize the difference in correct rates across odor difficulty to provide a relatively linear relationship between odor difficulty and correct trial rates. After the adjustments, the medium cue mixtures were typically defined as 56:44/44:56 and difficult cue mixtures as 51:49/49:51 with some variations as detailed in Table 1. Those adjustments of odors were made concurrently with (5).
7. In random subsets of trials within a session, the reward was withheld by setting the reward delay to 30 seconds (i.e., probe trials). We began with a low probability of probe trials and gradually increased it to 20%, which was used in the recording sessions. The probe trials aid the precise assessment of reward waiting time in correct trials, independent of the time spent on reward consumption. The introduction of the probe trial was made concurrently with (5) and (6)

### Behavioral data analysis

#### Decoding of session types from waiting time distributions

We trained a linear SVM to predict session types (variable long or fixed delay) based on the shape of the reward waiting time distribution. First, we computed the skewness and kurtosis of the reward waiting time distribution for each session. These values were then concatenated into a matrix with two parameters (skewness and kurtosis) × 134 sessions.

To eliminate any bias due to differences in the ranges of the parameters, each row of the matrix was normalized to the 0-1 range. The decoder was trained and tested using ten-fold cross-validation. A model cost function (C) was optimized to a value giving minimum loss of validation by searching in a range between 10^-5^ and 10^5^ using the training data^43^. To obtain a decoder performance distribution, this decoding procedure with cross validation was repeated 100 times. To determine chance level performance, another decoder was trained and tested with label shuffled data. The significance of the decoding performance was evaluated by comparing the real decoder performance distribution to the one obtained with label-shuffled data. Specifically, P-value was estimated as (1 + *X*) / (*N* + 1), where *X* represents the number of decoding performances below the chance level, and *N* represents the number of iterations. Since we repeated the decoding performance measurement 100 times, a decoder with no below chance performance resulted in *P* < 0.0099, and a distribution with *x*% overlap with chance level performance range resulted in *P* ≈ *x* /100^28,46^.

To obtain negative control, we also predicted the session types based on odor discrimination patterns. A data matrix comprising six conditions (left choice probabilities for six odor mixtures; Fig. 1B) × 134 sessions was constructed and used for decoding. The decoder was trained and tested following the same procedure described above, including the statistical test.

### Evaluation for metacognitive performance

To quantify performance in sensory discrimination, metacognitive (i.e., confidence) judgment, and metacognitive efficiency, we computed *d’*, meta-*d’*, and M-ratio (meta-*d’*/*d’*) using the MATLAB toolbox provided by Maniscalco & Lau (2012) (https://www.columbia.edu/∼bsm2105/type2sdt/)^40^. For each session, we first extracted only trials in which reward waiting time was measured (i.e., correct probe trials and all error trials). To ensure that the ratio of correct to error trials matched the true correct rate computed across all trials in the session, we randomly excluded a subset of error trials as needed. Odor cues were categorized into two stimulus types based on their associated correct choice directions (left or right), and the animal’s choice direction (left or right) was treated as the response category. Confidence ratings were divided into four levels by splitting the distribution of the reward waiting times within each session at the 25th, 50th, and 75th percentiles.

### Generalized linear model for behavior

Generalized linear models (GLMs) were employed to quantify the influence of various task factors on rats’ behavior. Models were fitted and tested for behavior prediction using five-fold cross-validation with the cvglmnet package for MATLAB (https://hastie.su.domains/glmnet_matlab/intro.html). Regularization was applied using a 95% L1 and 5% L2 penalty (Elastic Net). The hyperparameter λ was optimized by selecting the value that minimized the cross-validation loss (three-fold) within the training data at each iteration of the five-fold cross-validation. Model performance was evaluated using explained variance (R²) for Gaussian GLMs and binary prediction accuracy for logistic GLMs. To assess the influence of individual task variables on behavior, each factor was excluded from the full model, which included all factors, to create a partial model. The decline in performance (R² or accuracy) in the partial model was then quantified. Specifically, we calculated the relative contribution of each task factor by determining the percentage of the total performance decline attributed to each factor (citation). The partial model was created by refitting, which reruns model fitting without an excluded task variable.

1. Waiting time model: For each session, correct probe trials and all error trials were selected to analyze reward waiting times. A Gaussian GLM was used to predict trial-by-trial reward waiting times based on the following task variables: confidence (six levels, ranging from 0 to 1: easy error to easy correct), odor difficulty (three levels, ranging from 0 to 1: difficult to easy), previous trial outcome (0 or 1: unrewarded or rewarded), choice direction (-1 or 1: left or right), and reward satiety (0 to 1: accumulated reward amount across trials in a session).
2. Choice model: For each session, trials involving a left or right port choice were selected. A logistic GLM was used to predict trial-by-trial choice direction (left choice probability) based on the following task variables: odor cue (six types, ranging from 0 to 1), previous trial left reward (0 or 1: unrewarded or rewarded), previous trial right reward (0 or 1: unrewarded or rewarded), and reward satiety (ranging from 0 to 1). Model performance was assessed by computing prediction accuracy based on the left choice probability. A left choice probability greater than 0.5 was interpreted as the model predicting a left choice, while a probability less than or equal to 0.5 was interpreted as the model predicting a right choice. Prediction accuracy was defined as the proportion of predictions that correctly matched the rat’s actual choices.

### Surgery

Rats were anesthetized with 2.5% isoflurane before surgery, and it was maintained throughout surgical procedures. We monitored body temperature, movements, and hind leg reflex and adjusted the depth of the anesthesia as needed. Eye ointment was used to keep the eyes moistened throughout the surgery. Subcutaneous scalp injection of a lidocaine 1% solution provided local anesthesia before the incision. A craniotomy was made over the the left ventrolateral OFC (AP +3.0–4.6 mm, ML 2.3–2.4 mm relative to the bregma, 3.5–4.2 mm below the brain surface), and an array of electrodes was vertically implanted using a stereotactic manipulator. A stainless-steel screw was placed over the cerebellum and served as the ground during the recordings. Electrode placement was estimated based on the depth and was histologically confirmed at the end of the experiments (Supplementary Fig. 2A).

### Electrophysiological recording

We used shuttle drives or custom-designed micro-drives for the extracellular recordings (for two and five rats, respectively). Both design had 16 drivable tetrodes which consisted of four twisted 12.5 µm diameter tungsten wires (California Fine Wire, Grover Beach, CA) and signals are simultaneous obtained through 64 channels. The electrode typically had an impedance of 140–700 kΩ at 1 kHz. To aid visual inspection of electrode tracks after the experiment, we applied fluorescent dye (Vybrant™ DiI, Thermo Fisher, #V22885) to tetrodes. The signals were recorded with Open Ephys acquisition board (Open Ephys, Cambridge, MA) at a sampling rate of 30 kHz and bandpass filtered between 0.1 and 7.6 kHz. To facilitate comparison of neuronal activity between session types, electrode positions were kept constant across two consecutive sessions with different delay distributions. After completing recordings from such paired sessions, electrodes were advanced by 37.5 to 160 μm using screws on the drives.

### Histology

Once the experiments were completed, the rats were deeply anesthetized with sodium pentobarbital and then transcardially perfused with phosphate-buffered saline and 4% paraformaldehyde. The brains were removed and post-fixed in 4% paraformaldehyde, and 100 μm coronal sections of the brains were prepared to confirm the recording sites.

### Spike sorting

The recorded data from each session was first denoised by common average referencing, a preprocessing that subtracts the median signal of all channels from each channel’s signal to remove common artifacts. This was implemented with the Spikes package (https://github.com/cortex-lab/spikes). We then sorted the denoised signals into single units using Kilosort 2.0 (https://github.com/MouseLand/Kilosort) with the following adjusted ‘ops.’ parameters: fshigh = 600 (applied for two micro-drive animals), lambda = 16, and minFR = 0.01. The single units identified as ‘good’ clusters by Kilosort based on cross-correlogram contamination were then manually inspected and curated using Phy2.0 (https://github.com/cortex-lab/phy). Specifically, we first excluded units with physiologically unnatural features, classifying them as noise, by examining spike waveforms, signal localization across electrodes, and inter-spike interval patterns. We then merged survived clusters that were improperly split but belonged to the same unit and separated noise from units with apparent noise contamination. These curation processes resulted in an average of 11.6 well-isolated single units per recording session, with a total of 1553 units.

### Unit selection for GLM

We included all well-isolated single units in the neuronal activity analysis. However, for the GLM, the absence of firing activity during designated epochs (i.e., reward waiting time) resulted in ill-conditioned data. Consequently, only units with firing rates exceeding 1 Hz during the reward waiting time were included in the GLM-based analysis.

### Outcome prediction selectivity

To identify neurons encoding post-decision confidence, we performed receiver operating characteristic (ROC) analysis to quantify neuronal firing activity predictive of decision outcomes during the reward waiting time. For each neuron, we calculated the average firing rate during the reward waiting time for each trial. We then obtained the area under the ROC curve (AUC) from the average firing rates in correct probe trials and all error trials. Since confidence in this study is defined as the probability of choice correctness, the comparison between correct and error trials reflects the proximity of higher and lower confidence conditions^32^. To facilitate intuitive understanding of different selectivity types, outcome prediction selectivity was defined as 2×(AUC−0.5), ranging from −1 to 1, with zero indicating no selectivity^28,44^. In this analysis, a positive value represented increased neuronal firing for higher confidence (i.e., positive confidence neurons), whereas a negative value indicated the opposite pattern, with increased firing for lower confidence (i.e., negative confidence neurons). The statistical significance of the selectivity was assessed using a permutation test, which compared the selectivity derived from the true data to a distribution of selectivity values generated by randomly reassigning trials to the two conditions 1,000 times, irrespective of their true classes (correct/error trials). Neurons with permutation test *p*-values below 0.05 were considered to exhibit significant selectivity for post-decision confidence.

### Peri-event time histogram (PETH)

As the reward waiting time varied across trials, it was normalized to facilitate the visualization of neuronal activity in PETHs. Following previous studies that analyzed events with variable durations^22^, the interval between choice onset and offset in each trial was linearly scaled to match the median waiting time across all sessions, separately for each session type (long: 4.46 s; short: 1.25 s). During the waiting time in each trial, spike timings were used to calculate the mean firing rate in 10 ms bins, and a Gaussian kernel (σ = 50 ms) was applied to obtain smoothed PETHs. A time-scaling coefficient was calculated by dividing the waiting time of each trial by the median waiting time. The PETH time for each trial was then adjusted to the median waiting time by multiplying it by this coefficient. In cases where trial durations were extended beyond their original length, missing values were imputed using the median firing rate from the surrounding 10 bins. The time-normalized PETHs were then averaged across trials under the desired conditions to generate plots (Fig. 3). To avoid potential artifacts from normalization, all analyses focusing on the temporal dynamics of neuronal activity were performed without normalizing event durations (Fig. 4-6).

### GLMs for neuronal activity

Kernel-based GLMs were employed to quantify the influence of task variables, dependent on behavioral events, on the activity of individual neurons. Models were fitted and tested for behavior prediction using ten-fold cross-validation with the cvglmnet package for MATLAB. Given the partial correlation between confidence and odor difficulty, regularization was applied using a 95% L1 and 5% L2 penalty (Elastic Net). The hyperparameter *λ* was optimized by selecting the value that minimized the cross-validation loss (three-fold) within the training data at each iteration of the ten-fold cross-validation. Model performance was evaluated using explained variance (*R²*), and the influence of individual task factors was assessed by computing their relative contributions, following the same procedure as for the behavioral GLM. Partial models were generated by refitting the full model with one type of kernel excluded. Statistical significance of each kernel was determined by directly comparing full model *R²* with partial model *R²* using t-test with FDR correction (*α* = 0.025; Fig. 4C & J).

Instantaneous firing rates for each neuron on each trial were computed using 50 ms bins and then smoothed with a Gaussian kernel (*σ* = 150 ms). Because reward waiting time varied across trials, firing rates for model fitting were extracted from 0.1 seconds before the rat poked its snout into a port to 0.1 seconds after the rat left the chosen port (i.e., the waiting time offset). We included event-based Gaussian kernels (*σ* = 150 ms) representing combinations of three task events (waiting time onset, waiting time offset, and the interval between them) and three task variables (confidence, odor difficulty, and choice direction). A time-independent kernel representing trial-by-trial reward satiety was also included. In total, 10 different types of kernels were incorporated into the model (supplementary Fig. 3A).

### Clustering of neurons with similar task-related activity

To identify clusters of neurons with similar task-related activity patterns, a Gaussian Mixture Model (GMM) was applied to the matrix of relative contributions of 10 variables (i.e., combinations of task variables and events) obtained from the kernel-based GLM. Following a previous study^16,42^, the GMM was implemented with a diagonal covariance matrix constraint, a maximum of 10,000 iterations, a regularization value of 0.005, and 100 replicates to ensure reliable clustering. We only included neurons that were relatively better fitted (full model *R²* > 0.15; Fig. 4B), resulting in 353 and 433 neurons for the long and short delay sessions, respectively. GMM clustering was performed independently for the long and short delay sessions. The optimal number of clusters was determined by the Bayesian Information Criterion (BIC) score. BIC scores were computed for cluster numbers ranging from 1 to 20 (supplementary Fig. 3B), and the optimal cluster number determined by the minimum BIC score was selected in Fig.4 (supplementary Fig. 3C).

For visualization purposes, we further classified neurons within each cluster according to their tuning types (e.g., Fig. 4E). We computed trial-averaged instantaneous firing rates for each condition using 10 ms bins, smoothed with a Gaussian kernel (*σ* = 50 ms), from 0.3 seconds before to 1.8 seconds after the onset of the reward waiting time, and from 1.8 seconds before to 0.3 seconds after its offset. For averaging across neurons, the firing rates of individual neurons were then normalized to a 0–1 range. To determine confidence tuning types (positive or negative), we compared the time-averaged firing rates between correct and error trials. A neuron was classified as positively tuned to confidence if its average firing rate was higher in correct trials than in error trials. Conversely, neurons with higher firing rates in error trials were classified as negatively tuned (left panel in Fig. 4E). Odor difficulty tuning types were determined similarly by comparing firing rates between the easiest and most difficult odor trials. A neuron was classified as tuned to higher difficulty if its average firing rate was higher in the most difficult odor trials than in the lowest ones. Conversely, neurons with higher firing rates in the easiest odor trials were classified as tuned to easier difficulty (right panel in Fig. 4E).

### Population dynamics

To gain an intuitive understanding of the geometry of task representations in the OFC, we employed principal component analysis (PCA), an unsupervised dimensionality reduction technique widely used for analyzing population-level neural representations^20,28,43,46,48,49^.

For the analysis of population dynamics emerging from the neuron clusters identified by GMM (Fig. 4C & J), we constructed an N ^neurons^ × N ^time points ×conditions^ matrix for each session type. Since our primary interest was to capture population activity dynamics during the reward waiting time, we performed PCA on neural activity recorded from 0.1 s before to 1.8 s after the onset of the waiting period (Fig. 4H & N), and separately from 1.8 s before to 0.1 s after waiting time offset (Fig. 4I & O). Because the reward waiting time varied across trials, we excluded trials shorter than 1.8 s to ensure sufficient data range across time. Excluding trials with short waiting times often led to a low number of low-confidence trials per neuron. To minimize the influence of high variance from those conditions with small sample sizes, we excluded neurons that did not meet a minimum trial threshold (median number of trials across confidence levels < 5). After applying these criteria, we included 350 neurons for the long delay session and 209 neurons for the short delay session.

We analyzed five confidence levels (excluding easy error trials, which had high variance and low trial counts) and three odor difficulty levels as experimental conditions. This resulted in data matrices of size 350 × 1448 for the long delay session and 209 × 1448 for the short delay session. Prior to PCA, we z-scored each row to center and normalize the data. PCA was then applied separately to each session type. The resulting principal components were used to visualize population-level dynamics as trajectories in the low-dimensional PCA space (Fig. 4H–I & N–O).

### Time-resolved and cross-temporal population decoding

We trained a linear SVM to predict confidence levels from neuronal population activity in OFC. Since our interest was to analyze how decoding performance changed along the duration of the reward waiting time, we trained and tested SVM binary decoder using instantaneous firing rates of neurons (50 ms bins smoothed with Gaussian *σ* = 250 ms). To ensure sufficient time window for analysis, we excluded trials with waiting time shorter than

1.8 seconds. From the remaining data, we randomly selected 30 trials each from correct and error trials, corresponding to relatively high and low confidence levels, respectively. Neurons without enough trials to meet this criterion were excluded from the analysis. After this thresholding, 840 neurons remained for the long delay sessions and 343 for the short delay sessions. The firing rates of individual neurons were normalized to a 0–1 range. As a result, a data matrix of 840 or 343 neurons (long delay or short delay session) × 94 time bins (from 0.5 seconds before to 1.8 seconds after the waiting time onset, and from 1.8 seconds before to 0.5 seconds after the waiting time offset) × 60 trials was constructed for time-resolved decoding. To enable a direct comparison of decoding performance between session types, we randomly sampled neurons from the long delay session to match the population size of the short delay session. To minimize bias by neuron selection, random sampling was performed independently in each of the 100 decoding iterations (see below).

The decoder was trained and tested using five-fold cross-validation. A model cost function (C) was optimized to a value giving minimum loss of validation by searching in a range between 10^-5^ and 10^5^ using the training data. To obtain a decoder performance distribution, this decoding procedure with cross validation was repeated 100 times. The significance of the decoding performance was assessed by estimating *p*-values as described above, computed by comparing the distribution of real decoder performance with that obtained from a decoder trained with label-shuffled data (Fig. 6A & D).

To analyze how population coding patterns evolved through the duration of the reward waiting time, the decoders trained in the above time-resolved manner were also tested on the data at other time points. High performance of this cross-temporal decoding indicates that population coding patterns for confidence are consistent between different time points. We performed cross-temporal decoding for all combinations of time points around the waiting time onset (−0.5 to 1.8 s) and offset (−1.8 to 0.5 s), independently (Fig. 6B & E). To evaluate the temporal stability of population coding patterns, we compared the decoding performance at each off-diagonal time point (e.g., decoder trained at *t* = 0 s and tested at *t* = 1 s; cross-temporal decoding) with the corresponding on-diagonal time points (e.g., decoders trained and tested at *t* = 0 s and *t* = 1 s; within-time decoding), using estimated p- values as described above. Off-diagonal points were classified as reflecting *stable* population coding when decoding performance did not significantly drop relative to the relevant on-diagonal conditions (Fig. 6C & F).

To statistically compare the stability of population coding between the long and short delay sessions, we computed a stability index (Fig. 6G). For each time point, the stability index was defined as the proportion of relevant off-diagonal decoding performances that were not significantly lower than the 2.5th percentile of the corresponding on-diagonal decoding performances. Specifically, for each decoding iteration, if an off-diagonal performance exceeded this threshold, it was classified as stable. The proportion of stable off-diagonal performances across decoding iterations was then calculated for each time point and used as the stability index. Because the stability index was computed from off-diagonal data for each iteration, it resulted in a distribution across iterations. We compared the resulting distributions between the long and short delay sessions using estimated *p*-values.

### Representational geometry

For each session type, we constructed an N ^neurons^ × N ^conditions^ matrix (840 × 12 for long delay sessions; 713 × 12 for short delay sessions). The conditions consisted of six odor cue types and two choice outcomes (correct/error) which together can define different confidence levels. Each row of the matrix contained the time-averaged firing rates of a neuron during the reward waiting time across different conditions. To center and scale the data for PCA, we computed the z-score for each row. By performing PCA on these matrices, we reduced the dimensionality of the OFC population from 840 and 713 neurons to several principal components (PCs) (Fig. 7) .

### Generalization of population decoding

We trained a linear SVM to predict confidence levels from neuronal population activity in the OFC. To investigate the representational geometry of continuum of confidence levels at the population level, we took advantage of a decoder generalization approach, which is widely used to identify neuronal subspaces encoding abstract cognitive variables^16,23,25^.

We first selected neurons that had more than 20 trials for each confidence level, excluding the lowest level (i.e., easy error). To ensure reliable neuronal activity, we included only trials with reward waiting times longer than 0.5 seconds. After this thresholding, 658 neurons remained for the long delay session and 711 neurons for the short delay session. We then randomly selected 20 trials for each of five confidence levels: easy cue error + medium cue error, difficult cue error, difficult cue correct, medium cue correct, and easy cue correct. Because easy error trials were relatively rare, we supplemented the lowest confidence level by including medium error trials when fewer than 20 easy error trials were available. Next, we computed the time-averaged firing rate during the reward waiting time for each selected trial. The firing rates of individual neurons were normalized to a 0–1 range. This preprocessing resulted in a data matrix of 658 (long delay) or 711 (short delay) neurons × 100 trials for decoding. To enable a direct comparison of decoding performance between session types, we randomly sampled neurons from the short delay session to match the population size of the long delay session. To minimize sampling bias, this random selection was repeated independently for each of the 100 decoding iterations.

Similar to the time-resolved decoding described above, we trained and tested a decoder using ten-fold cross-validation, with the cost parameter C optimized by searching over a range from 10^-5^ to 10^5^. The training was performed with the two lowest confidence levels in the data matrix (i.e., easy error + medium error vs difficult error). It was then tested not only on the trained condition but also on higher confidence levels (i.e., distinguishing easy error + medium error from difficult correct, medium correct, or easy correct trials) (Fig. 8D-E). This decoder generalization approach evaluates how consistently confidence is encoded along a subspace defined by population activity. If the representation follows a linear axis aligned with the gradient of confidence levels, the generalization performance is expected to improve progressively as confidence increases (Fig. 8C). To obtain a decoder performance distribution, this decoding procedure with cross validation was repeated 100 times. The significance of the decoding performance was assessed by estimating *p*-values, computed by comparing the distribution of real decoder performance to that obtained from a decoder trained with label-shuffled data (Fig. 8D-E). One-vs-one decoding was also performed in the same manner as described above, except that the training and test data always came from the same pair of conditions (Fig. 8F).

To evaluate how the scale of the neuronal population influences confidence decoding, we repeated the decoder generalization test (using 5-fold cross-validation) with differently sized neuronal populations. Since our primary interest was in how successful decoder generalization in the long delay sessions was supported by population size, this analysis was restricted to data from the long delay sessions. Decoding was performed using population sizes ranging from 4 to 658 neurons, corresponding to 0.61% to 100% of the full population size (Fig. 8G). To minimize sampling bias, neurons were randomly sampled in each iteration, and decoding was repeated 200 times for each population size. The resulting decoding performance for each population size was analyzed in terms of general decoding performance and generalization ability (Fig. 8G–I). These were quantified by fitting a linear model to the decoded confidence levels, where the intercept (*β*₀) reflected the overall decoding performance and the slope (*β*_test_) represented generalization performance.

To more precisely relate behavioral performance to decoding performance, we performed similar decoding analyses at the single-session level using simultaneously recorded neurons (Fig. 9). We restricted the analysis to sessions in which five or more neurons were recorded simultaneously. This resulted in the inclusion of 40 long delay sessions and 45 short delay sessions. On average, 15.6 ± 1.0 and 14.6 ± 1.0 neurons were included per session in the long delay and short delay conditions, respectively.

## Acknowledgements

We would like to thank members of the Laboratory of Neural Information at Doshisha University for helpful discussion on the manuscript. This work was supported by JSPS KAKENHI (21K20702, 22K15632, 25K18958 to T.O.; 20H00109 to Y.S.; 23K23998 to J.H.) and the JST FOREST Program (JPMJFR204E to J.H.).

## Contributions

T.O., Y.S., and J.H. designed the experiments. T.O. and J.H. performed the experiments. T.O., Y.O., K.S., Y.T. and J.H. analyzed the data. H.M, Y.S., and J.H. supervised the project. All the authors contributed to writing and revising the manuscript.

## Corresponding authors

Correspondence to Tomoya Ohnuki and Junya Hirokawa.

---

**Supplementary Table 1.**
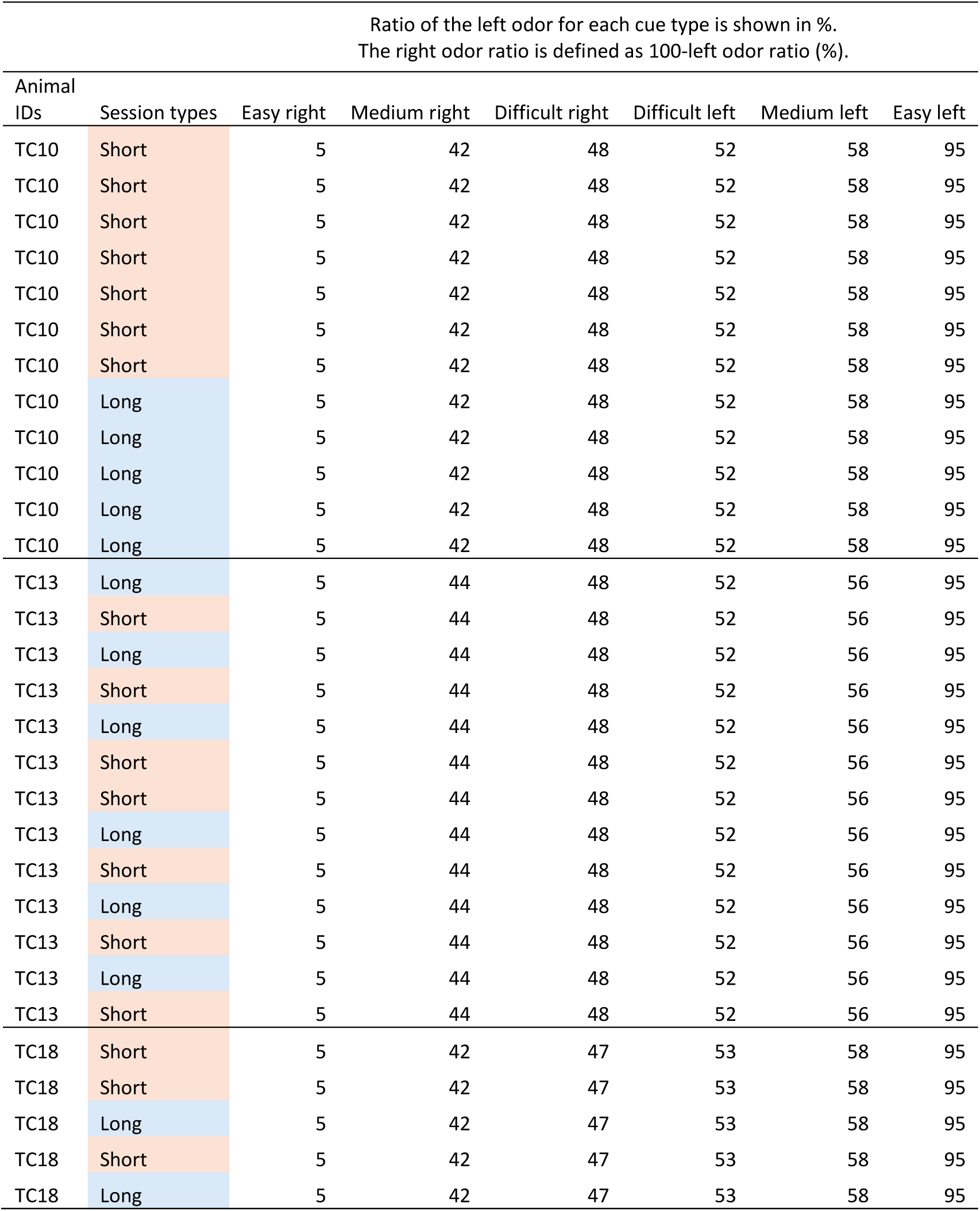
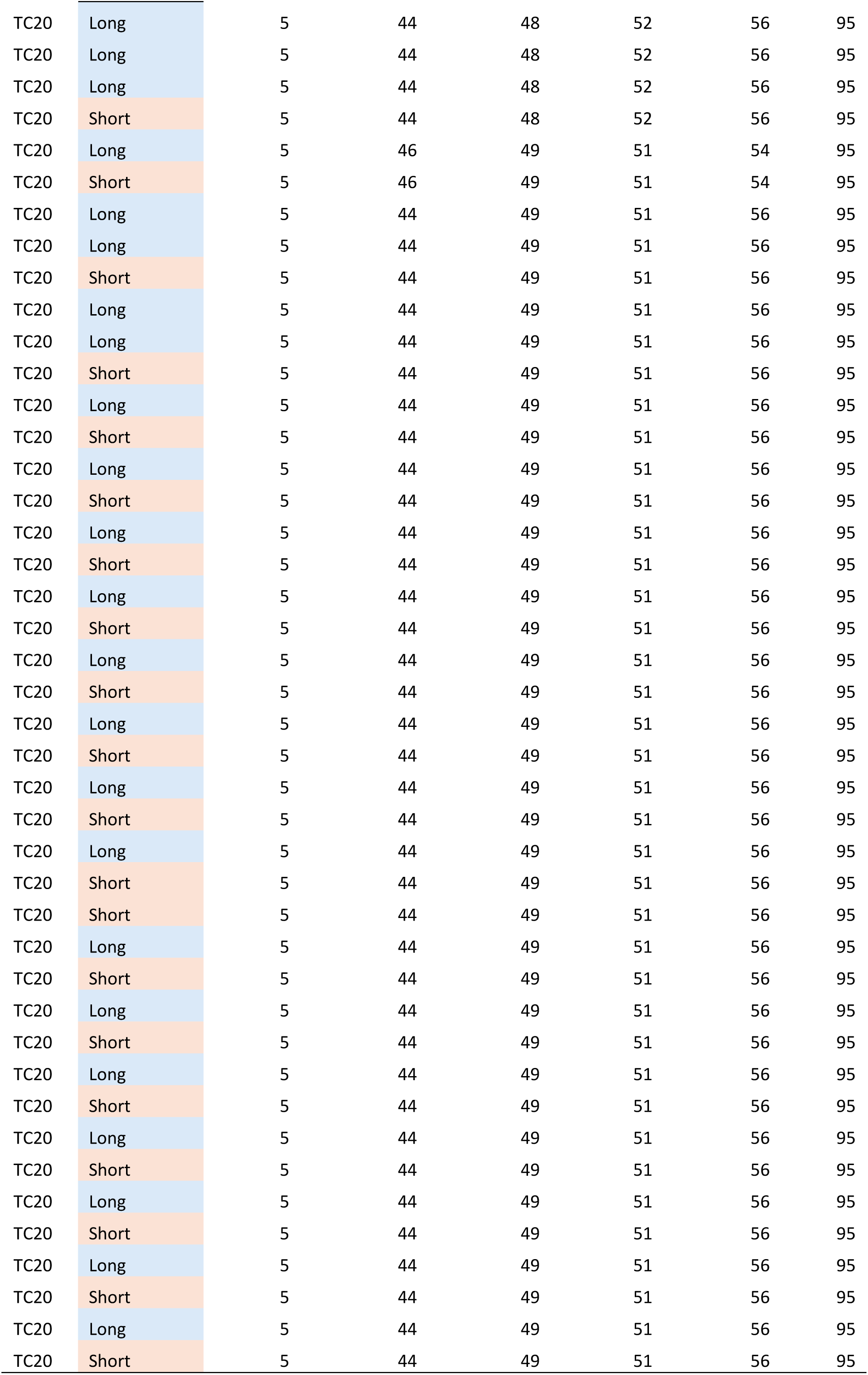
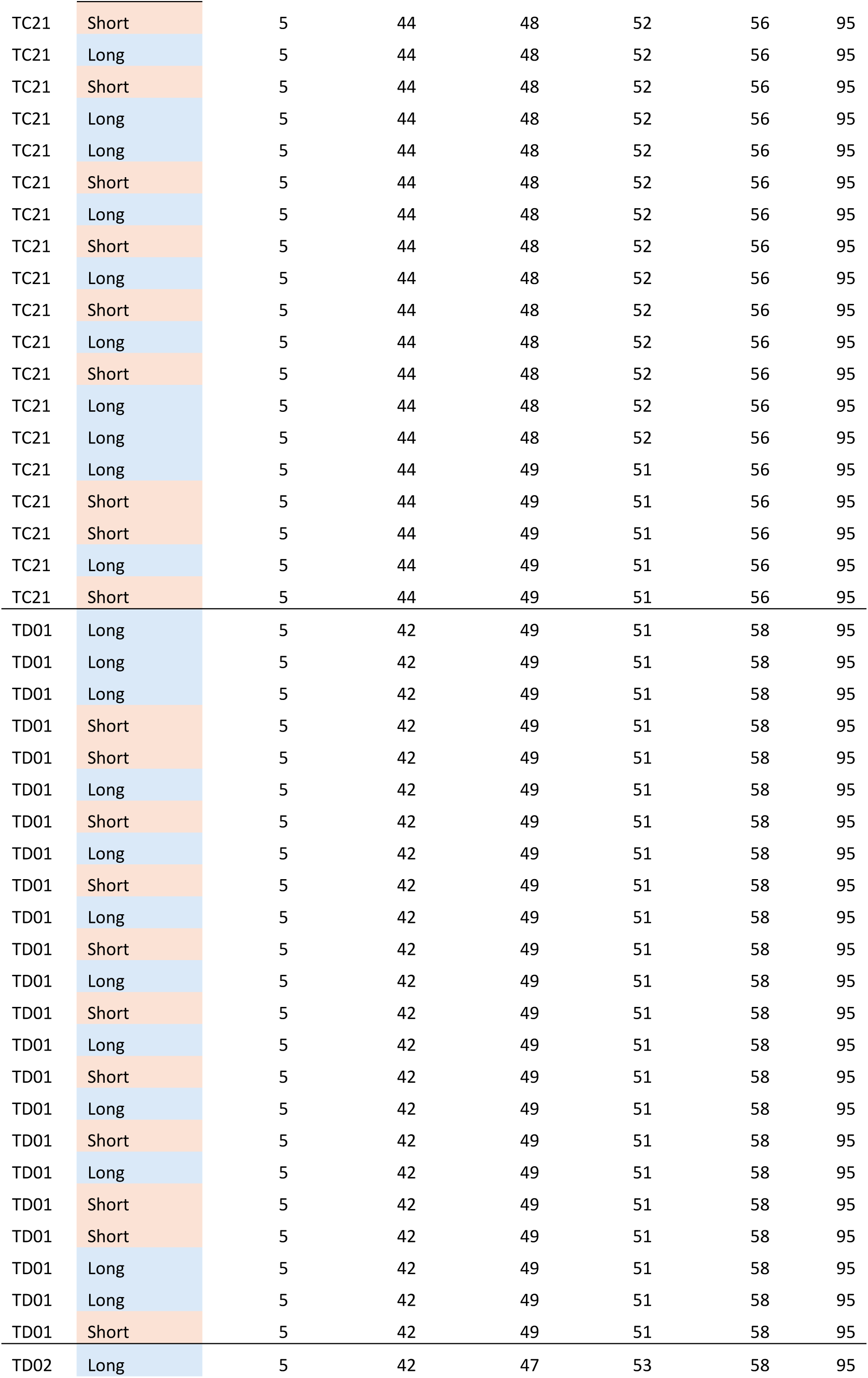
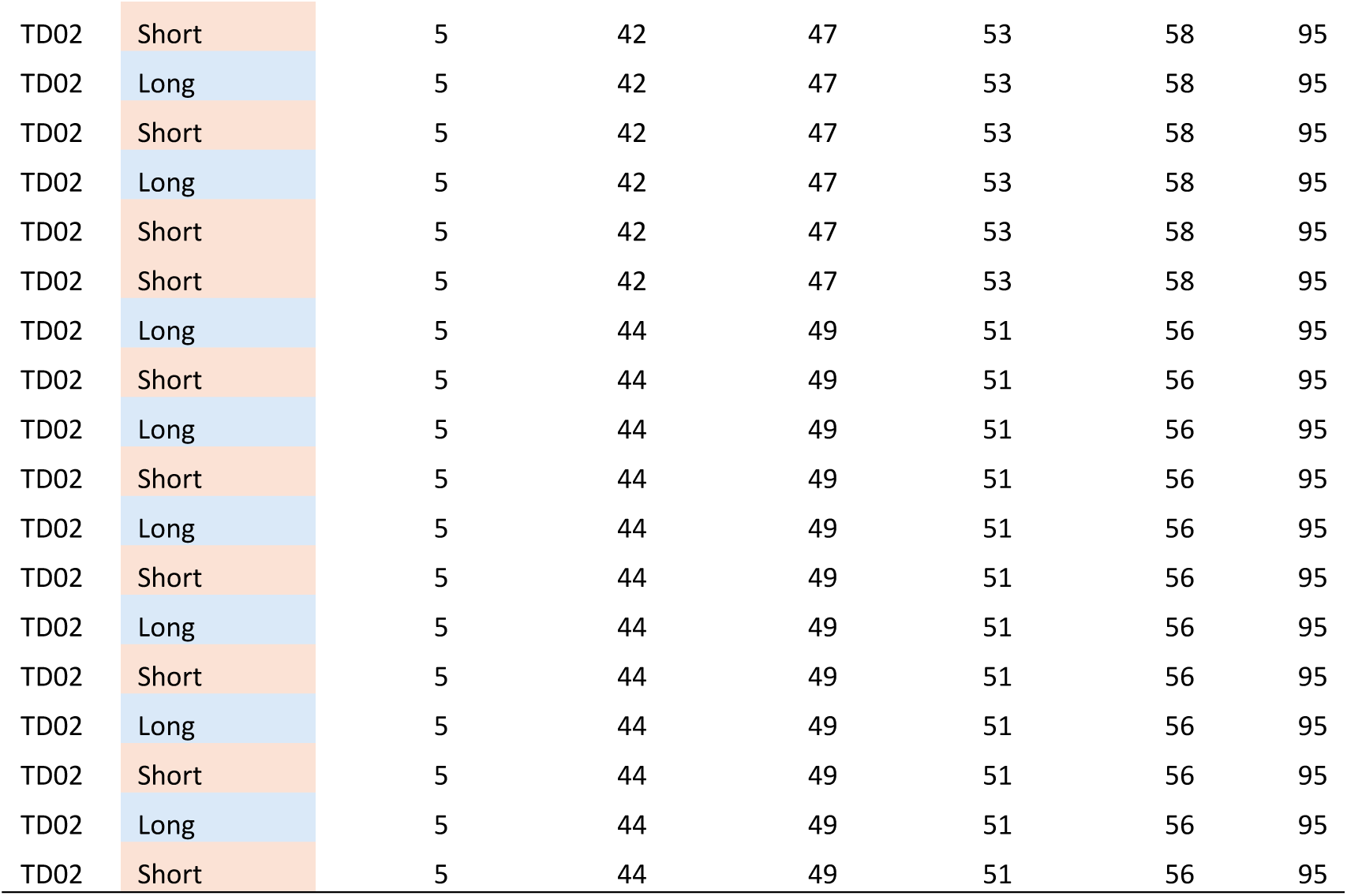
: Animals, session types, and odor mixtures for all recording sessions analyzed. Sessions in the table are presented in chronological order for each animal. For one rat (ID: TC10), recordings were conducted in a block design: data was recorded during a series of short delay sessions followed by a series of long delay sessions. For the remaining animals, the two session types were alternated across recording days. In some cases, the same session type appears in consecutive sessions because sessions without identifiable units were excluded from the analysis. Nonetheless, the most analyzed sessions maintained the intended alternation between the two types.

**Supplementary Fig. 1:**
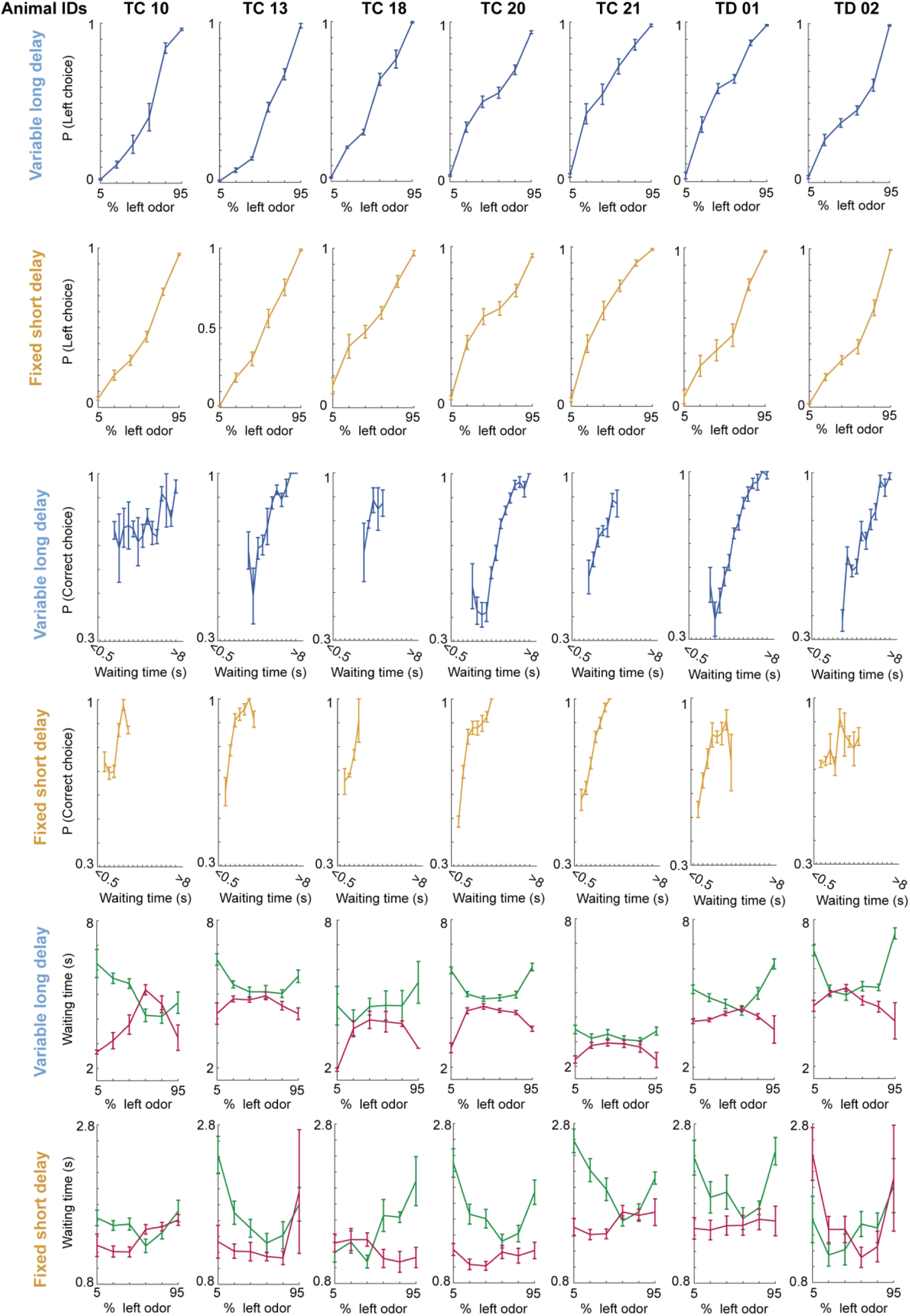
Individual animals show consistent trends across behavioral **measurements.** Each column corresponds to an individual animal, showing various behavioral measures presented in Fig. 1. The first and second rows show odor discrimination performance in the long delay and short delay sessions, respectively. The third and fourth rows display the correlation between reward waiting time and correct choice rate in the long and short delay sessions, respectively. The fifth and sixth rows present reward waiting time in the long and short delay sessions, respectively, conditioned on combinations of sensory evidence and choice outcome.

**Supplementary Fig. 2:**
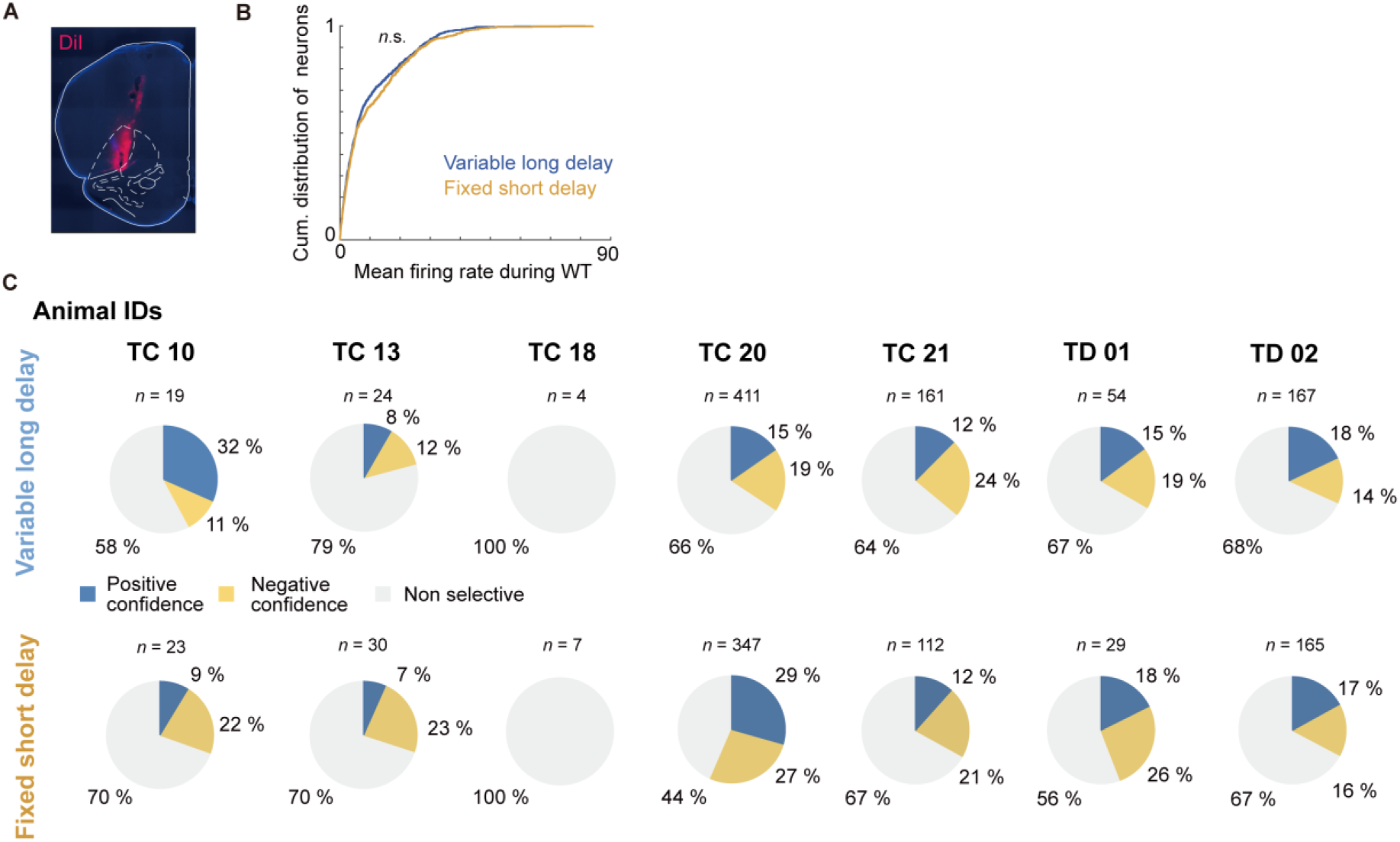
Neurons recorded from individual rats show consistent **trends in confidence encoding.** (A) Example of electrode placement in the ventrolateral OFC. (B) Comparison of time-averaged firing rates of all neurons during the reward waiting time between session types (Kolmogorov–Smirnov test, *p* = 0.115). (C) Proportion of neurons showing significant outcome prediction selectivity (permutation test; *p* < 0.05) during the reward waiting period. Each column represents an individual animal, and each row corresponds to a session type.

**Supplementary Fig. 3:**
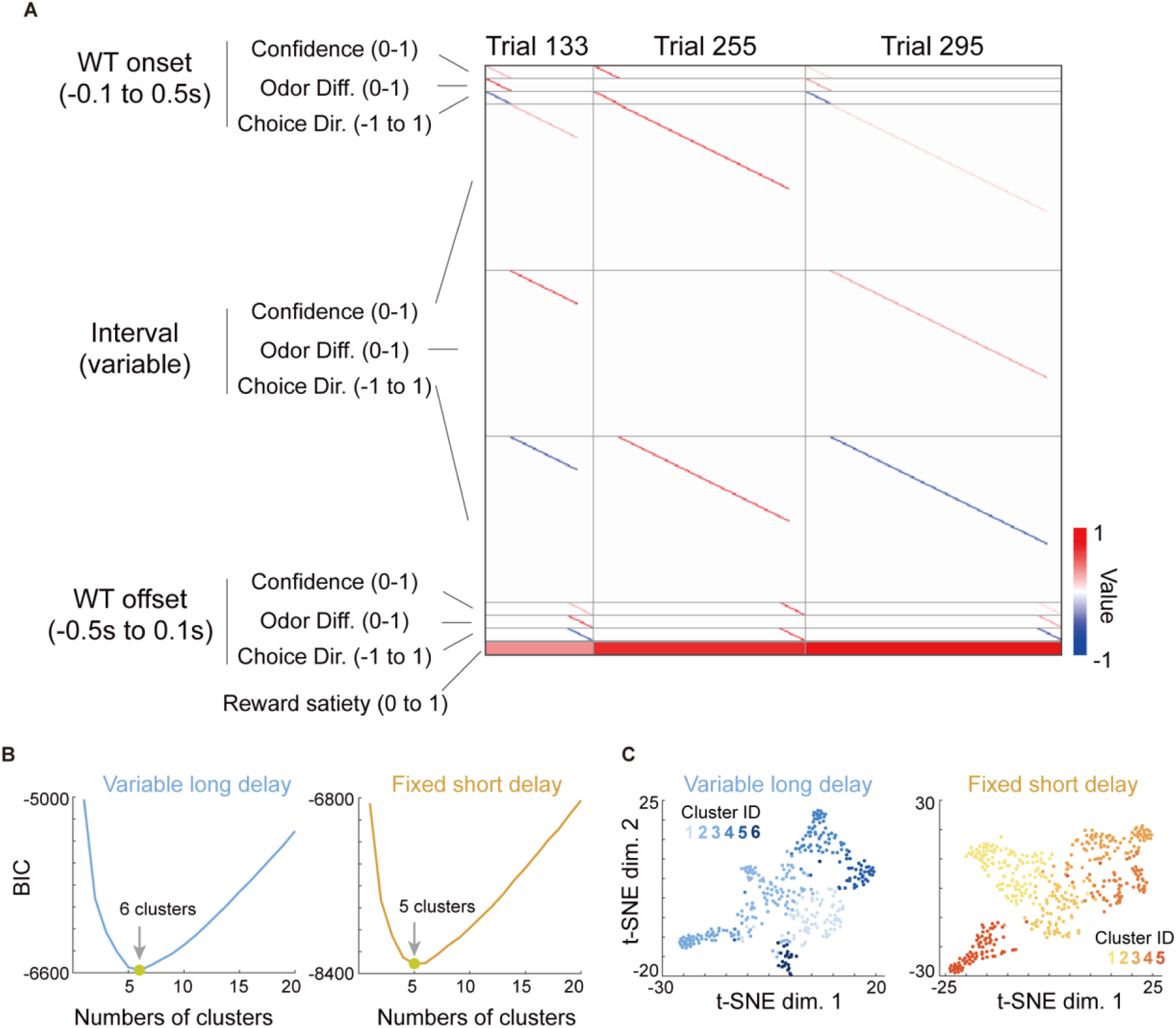
GLM analysis and GMM clustering. (A) Design matrix for the kernel-based GLM used to fit the instantaneous firing rates of individual neurons based on task variables and events. For illustration, three randomly sampled trials are concatenated and shown as columns. Each row represents a different type of kernel capturing either time-dependent or time-independent effects of task variables. (B) BIC values used in the GMM-based clustering analysis in Fig. 4. Arrows indicate the number of clusters that minimized the BIC. (C) Visualization of functional clusters of neurons identified through GMM-based clustering. The same clusters are shown in Fig. 4. t-SNE here was used solely for visualizing the high-dimensional data in two dimensions.

